# Foliar striping and floral traits are correlated with higher reproductive output in crimson monkeyflowers

**DOI:** 10.64898/2026.09.10.750677

**Authors:** Erika R. LaPlante, Liza M. Holeski

## Abstract

In plants, ecological interactions such as pollination depend on whole-plant phenotypes, where coordinated variation among vegetative, floral, and reproductive traits can shape reproductive success. However, empirical studies linking integrated phenotypes that include non-floral traits to reproductive outcomes within populations remain limited.

While prior work has shown that crimson monkeyflowers exhibit heritable variation in foliar anthocyanin striping, the functional significance of the trait has been thus far unknown. Using plants derived from natural populations, we conducted field and greenhouse experiments to assess whether foliar striping is genetically correlated with traits related to pollinator attraction. We found that foliar striped plants consistently exhibit larger flowers, greater nectar rewards, and decreased proportions of viable pollen relative to non-striped plants. When we manipulated flowers within plants with and without foliar striping to allow only outcrossing or only self-fertilization in a field environment, we found that outcrossed flowers had higher reproductive success than did self-fertilized, with outcrossed flowers from plants with foliar striping producing the highest overall seed numbers and seed mass.

Our results reveal patterns of coordinated foliar and floral trait variation and associated reproductive output that are consistent with a pollination syndrome framework. The link between foliar pigmentation and increased per-calyx reproductive output represents a novel association between foliar pigmentation and a broader evolutionary and ecological function.

## INTRODUCTION

In plants, ecological interactions such as pollination occur at the level of whole-organism phenotypes; coordinated variation among vegetative, floral, and reproductive traits may jointly influence reproductive success (Campbell 1989; Fenster 1991; Song et al. 2018; Leimberger et al. 2022; Young 1982). Multivariate selection theory provides a framework for understanding how suites of correlated traits can shape fitness and natural selection (Phillips and Arnold 1989; Jablonski 2008; Damián et al. 2020), yet empirical tests linking integrated suites of traits to reproductive outcomes within populations remain comparatively rare (Armbruster et al. 2014; Pigliucci 2003; Walsh and Blows 2009). One framework for conceptualizing trait integration is the pollination syndrome paradigm, which links suites of traits to predictable pollinator interactions (Leimberger et al. 2022; Campbell 1996; Simpson 1944; Wright 1931; Fenster et al. 2004).

Pollination syndromes link suites of plant traits (e.g. floral morphology, color, scent, and reward type) to particular pollinator functional groups; these have historically emphasized floral traits over vegetative or whole-plant signals (Faegri and Van der Pijl’s 2013; Dellinger 2020; Sinnott-Armstrong et al. 2022). Classic studies of hummingbird pollinated plants demonstrate strong pollinator-mediated selection on floral traits that enhance detectability and access, including red coloration, wide corollas, stigma exsertion, and relatively high nectar volume production (Campbell 1989, 1996; Schemske and Bradshaw 1999). Empirical studies demonstrate that hummingbirds have highly developed color vision and use visual cues when locating floral resources (Altshuler 2003; Hurly and Healy 1996; Stoddard et al. 2020). Hummingbirds integrate signals at spatial scales beyond individual flowers (such as vegetation density, background color, and canopy context) when selecting foraging sites, which indicates that cues provided by leaves and plant structure can shape pollinator behavior independently of floral traits (Meléndez-Ackerman et al. 1997; Altshuler and Dudley 2002). These findings suggest that pollinator visitation and effectiveness in hummingbird-pollinated systems may not be fully explained by floral traits alone and that foliar traits are likely an understudied component of pollination syndromes.

Foliar anthocyanins may function as visual signals that contribute to whole-plant display, particularly for hummingbirds that often detect plants from a distance (Lisney, Kolominsky, and Iwaniuk 2015; Fellows 2015; Altshuler and Wylie 2020). Here, we investigate the ecological role of a unique foliar anthocyanin striping trait in crimson monkeyflower *(Mimulus verbenaceus* Greene (1885) syn. *Erythranthe verbenacea* Greene (1909); (Nesom 2014)), including how it may affect plant-pollinator interactions. Some crimson monkeyflower plants express a lateral red/purple stripe across otherwise green leaves, caused by localized anthocyanin accumulation. This foliar striping phenotype is a relatively recently evolved trait that has risen to high frequency in some natural populations (LaPlante and Holeski personal observation; Weiss, Faske, and Holeski 2026). Previous work has identified the causal gene underlying this pattern, STRIPY, which encodes an anthocyanin-activating R2R3-MYB transcription factor (Yuan et al. 2014; LaFountain et al. 2023). Although the genetic and developmental basis of this foliar striping pattern has been well characterized, the ecological and evolutionary processes through which this phenotype has risen in frequency in natural populations remain unresolved.

Most research on anthocyanins in plant-animal interactions has focused on floral tissues, where color variation plays a central role in pollinator attraction. Anthocyanin-based red floral coloration is a hallmark of bird pollination syndromes and has been shown experimentally to influence pollinator visitation patterns, including shifts toward hummingbird pollination and away from bee visitation (Schemske and Bradshaw 1999). Although anthocyanins are also widespread in vegetative tissues, research on foliar pigmentation has largely emphasized physiological and defensive roles rather than pollination ecology. Foliar anthocyanins are well documented as photoprotectants, antioxidants, and stress-response compounds in vegetative tissues (Chalker-Scott 1999; Lev-Yadun et al. 2002). Vegetative tissues have also been implicated in herbivore deterrence through visual or chemical signaling of reduced palatability or elevated defense investment (Lev-Yadun et al. 2002; Lev-Yadun and Gould 2008; Ahmed et al. 2014). In the rare instances where leaf pigmentation and patterns have been considered in the context of pollination, they have been found to enhance floral display contrast or contribute to ultraviolet patterning (Benzing and Friedman 1981; Koski et al. 2024).

The crimson monkeyflower foliar striping system, while well-characterized genetically, is understudied from an evolutionary and ecological perspective. This system offers a rare opportunity to characterize how foliar pigmentation integrates with other traits to influence pollination and plant reproduction in natural populations. To do this, we address three questions via a greenhouse and a field experiment: [1] **Could foliar striping be part of a larger pollination syndrome** (i.e., is it genetically correlated with traits known to affect pollinator attraction)? [2] (a) **Is foliar striping associated with increased rates of successful pollination, if so, does this** (b) **result in higher relative reproductive output in foliar striped plants?**

## METHODS

### Materials

Crimson monkeyflower is an herbaceous perennial found in riparian habitats across northwestern Mexico and southwestern U.S. (Figure 1A). Plants typically occur along streams, rivers, and hanging gardens, overwintering as basal rosettes (iNaturalist 2024; Vickery 2008). The species produces red tubular flowers that are primarily pollinated by hummingbirds (Yuan 2019; Vickery 2008; Vickery and Vickery 1992). Crimson monkeyflower is also capable of self-fertilization, facilitated by close proximity of the stigma and anthers following anthesis, as well as by a delayed selfing mechanism in which the anthers attached to the senescing corolla contact the stigma (Vickery 2008).

**Figure 1.**
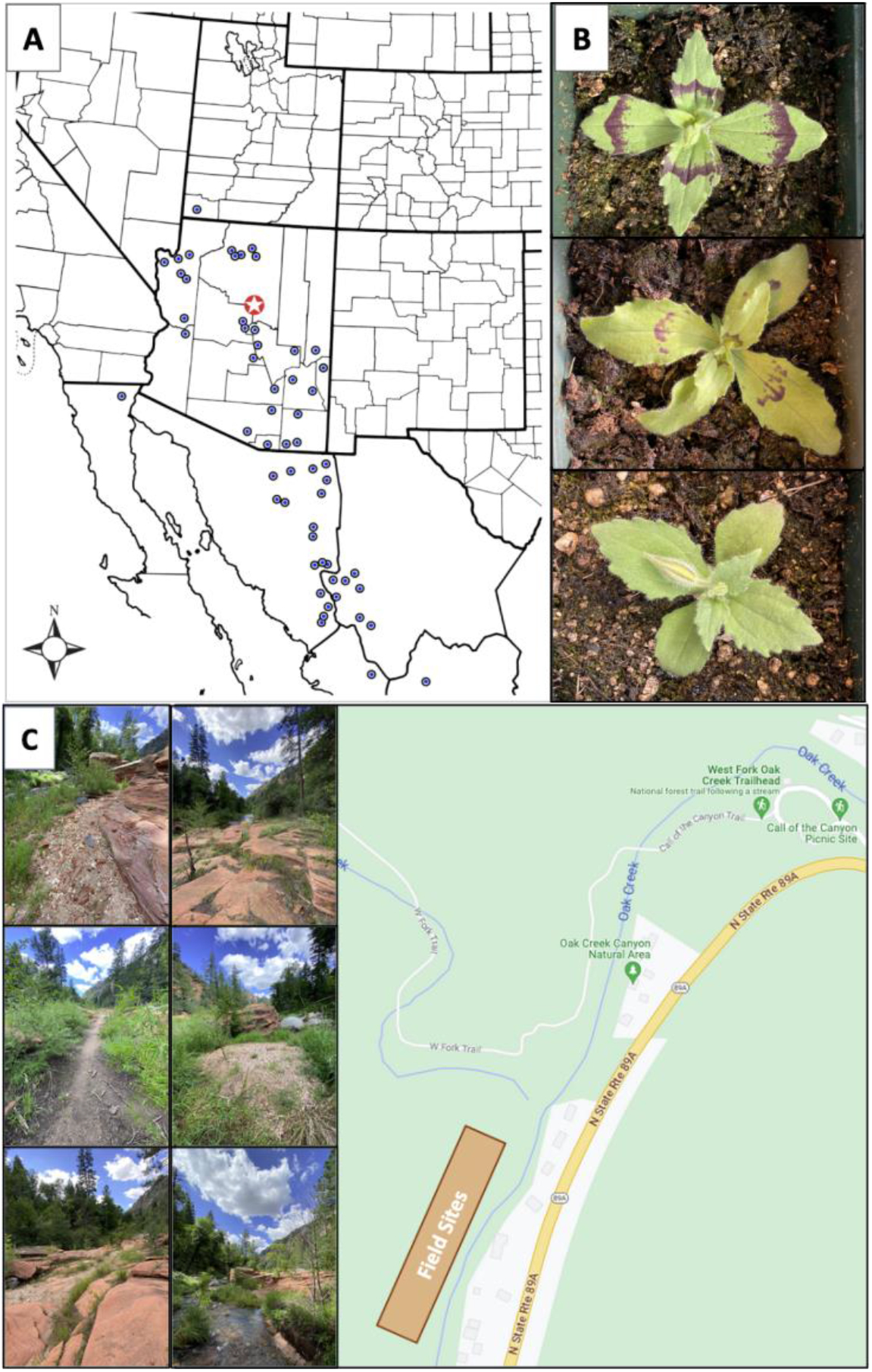
(**A**) Geographic distribution of natural crimson monkeyflower populations (blue dots). Experimental field sites and source populations are indicated by the red dot with a star, in panel A. (**B**) All three phenotypes seen in natural populations (intermixed in locations like West Fork of Oak Creek Canyon in Sedona, AZ) are also found in F_2_ individuals. (**C**) Experimental field site at West Fork of Oak Creek Canyon. The photographs on the left show the expanse of where the plant arrays were spread out across the site. The field site in relation to West Fork and Oak Creek Canyon is shown on the right.

In monkeyflowers, research on pigmentation has focused largely on floral anthocyanins. In multiple monkeyflower species, variation in floral color and patterning is thought to affect pollinator-mediated reproductive isolation (Liang et al. 2022; Peng et al. 2017; Chen et al. 2025; Lowry et al. 2012; Ding et al. 2020; Yuan et al. 2013, 2014, 2016; Yuan 2019) though few studies have shown a direct effect on the recruitment of pollinators and the resulting increased fitness of the plants (Schemske and Bradshaw 1999; Wenzell et al. 2025). In contrast, the ecological and evolutionary significance of anthocyanin pigmentation in leaves is comparatively unexplored. Existing work has focused on the molecular genetics of the phenotype, examining the genetic basis of phenotypic variation in floral and foliar tissues in yellow monkeyflowers and the repression of anthocyanin biosynthesis in the leaves of crimson monkeyflowers (Lowry et al. 2012; LaFountain and Yuan 2021).

To address our questions, we conducted a greenhouse and a field experiment using crimson monkeyflower plants with and without foliar striping. For the greenhouse experiment, we used parental lines with foliar striped and non-striped phenotypes, respectively, as well as an F_2_ family derived from a cross between a foliar striped and a non-striped parent plant, followed by self-fertilization of an F_1_ offspring. In these lines, foliar striping is controlled by a single locus (previously characterized as STRIPY) with a semi-dominant allele. F_1_ heterozygotes exhibit a faint stripe, often appearing as horizontally aligned spots in the same position as the full stripe, while F_2_ families segregate approximately 1:2:1 (strong stripe: faint stripe: no stripe; 86:191:116), consistent with Mendelian inheritance of a single locus (Figure 1B; LaFountain et al. 2023).

For the field experiment, we derived new experimental lines from seed collected from natural populations at the West Fork of Oak Creek Canyon (Sedona, AZ) (Figure 1C). Seeds were collected from naturally occurring striped and non-striped plants, grown in the greenhouse, and self-fertilized to generate inbred parental lines. A foliar striped and a non-striped parent were crossed to produce F_1_ individuals, one of which was subsequently selfed to generate an F_2_ family. As in the greenhouse experiments, all three phenotypes observed in F_2_ families occur naturally across the species range.

### Greenhouse Experiment

#### Floral Traits

To characterize genetic-based patterns of trait variation in the absence of natural pollinators, we quantified floral and reproductive traits hypothesized to be associated with pollinator recruitment and pollen transfer, including corolla width, corolla height, stigma-anther separation (herkogamy), nectar volume, and pollen viability in foliar striped and non-striped inbred parental lines (corolla width, corolla height, herkogamy: n=514; nectar volume: n=81; pollen viability: n=85), as well as F_2_ individuals (corolla width, corolla height, herkogamy: n=401; nectar volume n=108; pollen viability n=68) (Supplemental Table 1). We took measurements within the first 24 hours of anthesis for each of the first five flowers produced per plant. We measured corolla width (mm) at the widest point across the lower petals, and corolla height (mm) was measured from the base of the receptacle to the tallest petal.

In many flowering plants, like the crimson monkeyflowers, floral organs continue to reposition around anthesis, altering stigma-anther spatial relationships during the flower’s receptive period (Armbruster et al. 2014; Lloyd and Webb 1986; Webb and Lloyd 1986; Ye et al. 2019), which makes early measurement essential for capturing functionally relevant separation prior to pollination. Stigma-anther separation values ≤0, in which stigmas are positioned at or above anthers (approach herkogamy), reduce opportunities for autonomous selfing and are commonly associated with increased likelihood of outcrossing through pollinator mediation (Barrett 2003; Lázaro, Seguí and Santamaría 2020). Stigma-anther separation values > 0 indicate that the stigmas are below the anthers (reverse herkogamy), which carries a greater likelihood of self-fertilization. In our statistical analysis, we also coded herkogamy as a binary factor, with reverse herkogamy (positive stigma-anther separation) compared against approach or no herkogamy (zero or negative separation).

Nectar volume acts as a floral reward, influencing pollinator attraction and visitation rates, whereas pollen viability is a critical component of male reproductive success by determining the potential for successful pollen export and fertilization (Kessler and Baldwin 2015; Minnaar et al. 2018). For a subset of plants, we measured nectar volume using a 5-µL microcapillary tube inserted into the floral receptacle at the nectar gland. After 30 seconds, we measured the length of nectar in the tube (mm) and converted it to volume using the equation (V=πr^2^h), where r = 0.15 mm (half the internal diameter of the capillary tube) and (h) is nectar column length.

On another subset of plants, we measured proportional pollen viability per flower using lactophenol cotton blue staining (Carr and Dudash 1997; Kearns and Inouye 1993). We suspended pollen from a single flower in stain, and two 10-µL aliquots were pipetted onto a microscope slide and covered with a coverslip. We examined slides under 100x magnification, and pollen grains were counted using a predefined serpentine scanning pattern. We scored pollen grains as viable or inviable based on size and staining intensity, with large dark-blue stained grains scored as viable and smaller lightly stained grains scored as inviable (Carr and Dudash 1997; Kearns and Inouye 1993). We modeled pollen viability as a binomial response (viable vs. inviable grains), testing whether phenotype, genotype, and their interaction affected the proportion of viable pollen grains.

#### Per-Calyx Reproductive Output

To assess reproductive output when pollen is not limiting, we quantified seed production following controlled hand-pollinations in the greenhouse where stigmas were saturated with pollen. This approach allowed us to evaluate whether foliar striping was associated with genetic-based differences in reproductive capacity independent of pollinator visitation or efficiency of pollen transfer. For each of 164 plants, we manually self-fertilized five flowers and manually outcrossed five flowers using pollen from designated donor plants, resulting in ten experimental flowers per individual. We tracked mating system treatment and pollen donor identity using distinct floral markers. After seeds matured, we collected calyxes and stored them in individually labeled coin envelopes prior to processing. For each calyx, we quantified multiple aspects of the seed set: (1) seed number, (2) seed mass, and (3) average seed mass per individual seed. These metrics provided measures of reproductive output under standardized pollination conditions, allowing later comparison with field-based fitness patterns where reproductive success may vary due to pollinator or plant behavior.

### Field Experiment

To quantify trait variation and reproductive outcomes in the presence of natural pollinators while controlling mating context at the floral level, we conducted a field experiment at the West Fork of Oak Creek Canyon (Figure 1C). The experimental population consisted of 182 foliar striped parental plants, 178 non-striped parental plants, and 120 F_2_ family plants (23 striped and 97 non-striped). Plants remained in individual pots that were held in bottom-watering trays and were arranged in eight spatial arrays distributed across the field site. Arrays varied in phenotypic composition and included foliar striped-only parental arrays, non-striped-only parental arrays, mixed parental arrays, and mixed F_2_ family arrays.

#### Floral Traits

As in the greenhouse experiment, we measured multiple floral traits to assess the relationship between foliar striping and floral traits hypothesized to be associated with pollinator recruitment and pollen transfer. Within the first 24 hours of anthesis we measured corolla width, corolla height, and stigma-anther separation (herkogamy) in four flowers per plant in both striped and non-striped parental lines (n=1,318), as well as F_2_ individuals (n=491) (Supplemental Table 1). We did not assess nectar volume and pollen viability in this experiment.

#### Reproductive Success

To evaluate whether foliar striping was associated with differences in pollination success and subsequent reproductive output, we quantified multiple components of reproduction under controlled mating system. For each plant, we assigned the same four flowers used for floral trait measurements to experimental mating system treatments. Immediately after we took floral trait measurements, we haphazardly assigned two flowers per plant to obligately self-fertilize by enclosing unpollinated flowers in mesh bags, and two flowers to obligately outcross via pollination by emasculating flowers immediately after anthesis.

We collected calyxes after seed maturation and processed them to quantify four reproductive fitness metrics: (1) presence or absence of seed set, (2) seed number per calyx, amongst flowers that produced seed (3) seed mass per calyx, and (4) average seed mass per individual seed. These measures allowed us to evaluate presence or absence of any reproductive success in flowers reliant on either self-fertilization or external pollination, as well as the extent to which each mating system was successful if at least one seed was produced.

### Statistical Analyses

We conducted all analyses in R (version 2025.09.0+387; R Core Team 2023), using functions from the lme4 (Bates et al. 2015), lmerTest, emmeans (Lenth 2023), and DHARMa (Hartig and Hartig 2017) packages, with additional packages for data manipulation and visualization. We analyzed greenhouse and field datasets separately due to differences in experimental design, methods of pollination, and sources of non-independence. Sample sizes varied among traits in the greenhouse experiments because of differences in flowering, nectar production, and pollen availability (see Supplemental Table 1).

#### Correlation Analyses

To assess pairwise associations among floral and reproductive traits, we calculated Spearman rank correlation coefficients (ρ) separately by foliar phenotype (striped vs. non-striped) and, where appropriate, by mating system (selfed vs. outcrossed). We used Spearman correlations due to non-normal trait value distributions. Correlation matrices display all pairwise trait combinations with corresponding ρ values, and we evaluated significance using false discovery rate (FDR)-adjusted p-values to account for multiple comparisons.

#### Principal Components Analysis

We conducted principal components analyses (PCA) on standardized floral and reproductive fitness-related traits to evaluate multivariate trait covariation. We analyzed PC1 and PC2 scores using linear models to test for effects of Phenotype and Genotype.

#### Greenhouse Experiment

To evaluate effects of foliar phenotype, genotype (parent lines vs. F_2_), and mating system (selfed vs. outcrossed) under greenhouse conditions, we fit LMMs, GLMMs, or linear models (LMs) as appropriate for each response variable. For floral traits, we selected models based on error structure. We analyzed corolla width and corolla height using LMMs with Phenotype and Genotype as fixed effects and Plant ID nested within Experiment as a random effect. We analyzed nectar volume using a linear model with Phenotype and Genotype as fixed effects. We analyzed binary herkogamy (approach vs. reverse) using a binomial GLMM with a logit link. We analyzed pollen viability using a binomial generalized linear model with viable and inviable pollen counts modeled as a two-column response.

We analyzed seed number and average seed mass per calyx using LMMs with Phenotype, Genotype, Mating System Treatment, and their interactions as fixed effects and Plant ID as a random intercept to account for repeated measures across flowers. We analyzed average individual seed mass using a linear model with the same fixed effects structure.

#### Field Experiment

We analyzed field floral traits using LMMs or binomial GLMMs, depending on error structure, with Phenotype, Genotype, and their interaction as fixed effects and Plant ID nested within Array as a random effect. We analyzed seed production in two stages. First, we modeled seed presence (0 = no seeds, 1 = seeds present) using binomial GLMMs with a logit link to test whether foliar striping and mating context influenced pollination success. Models included Phenotype (striped vs. non-striped), Genotype (parent line vs. F_2_), Mating System Treatment, and, where appropriate, the Phenotype × Mating System Treatment interaction as fixed effects. We included Array and Plant ID as random intercepts.

Second, among flowers that produced at least one seed, we analyzed seed number and seed mass using linear mixed-effects models (LMMs) with Gaussian errors, as alternative distributions did not improve model fit. These models included Phenotype, Genotype, Mating System Treatment, and their interactions as fixed effects, with Plant ID nested within Array as a random effect.

#### Model Evaluation and Post Hoc Tests

We evaluated fixed effects in mixed models using Wald χ² tests derived from model summaries or Type II/III analyses of variance, as appropriate. We assessed model assumptions through visual inspection of residuals for Gaussian models and simulation-based diagnostics for generalized models using the DHARMa package. When assumptions for parametric inference were not met, we conducted pairwise Wilcoxon tests for post hoc comparisons. When alternative model structures did not adequately resolve skewness, we retained variables on their original scale or analyzed them as biologically meaningful binary responses rather than applying transformations. In the results section, all reporting of phenotype, genotype, and treatment means and corresponding differences are calculated from estimated marginal means that were obtained from the fitted models.

## RESULTS

**(Q1) Could foliar striping be part of a larger pollination syndrome (i.e., is it genetically correlated with traits known to affect pollinator attraction)?**

We focus on our field experiment results for traits measured in both the greenhouse and field (corolla width, corolla height, herkogamy), with greenhouse results referenced in our supplementary data. Nectar volume and pollen viability were measured only in the greenhouse.

### Corolla Width and Corolla Height

In the field experiment, foliar anthocyanin striping was associated with both increased corolla width and height (Table 1A, 2A; Figure 2A, B). Corolla width was significantly associated with the foliar striping phenotype, genotype (parental line vs. F_2_), and their interaction, with striped plants producing corollas that were 1.06x wider than corollas of non-striped plants. Parent plants produced corollas smaller than the F_2_ plants, with parent plant corollas an average of 0.93x the corolla width of F_2_ plants. The significant phenotype x genotype interaction indicates that the magnitude of the relationship between foliar striping and corolla width differed between parental lines and F_2_ families, with a larger difference in corolla width between foliar striped vs. non-striped F_2_ than in parents.

**Figure 2.**
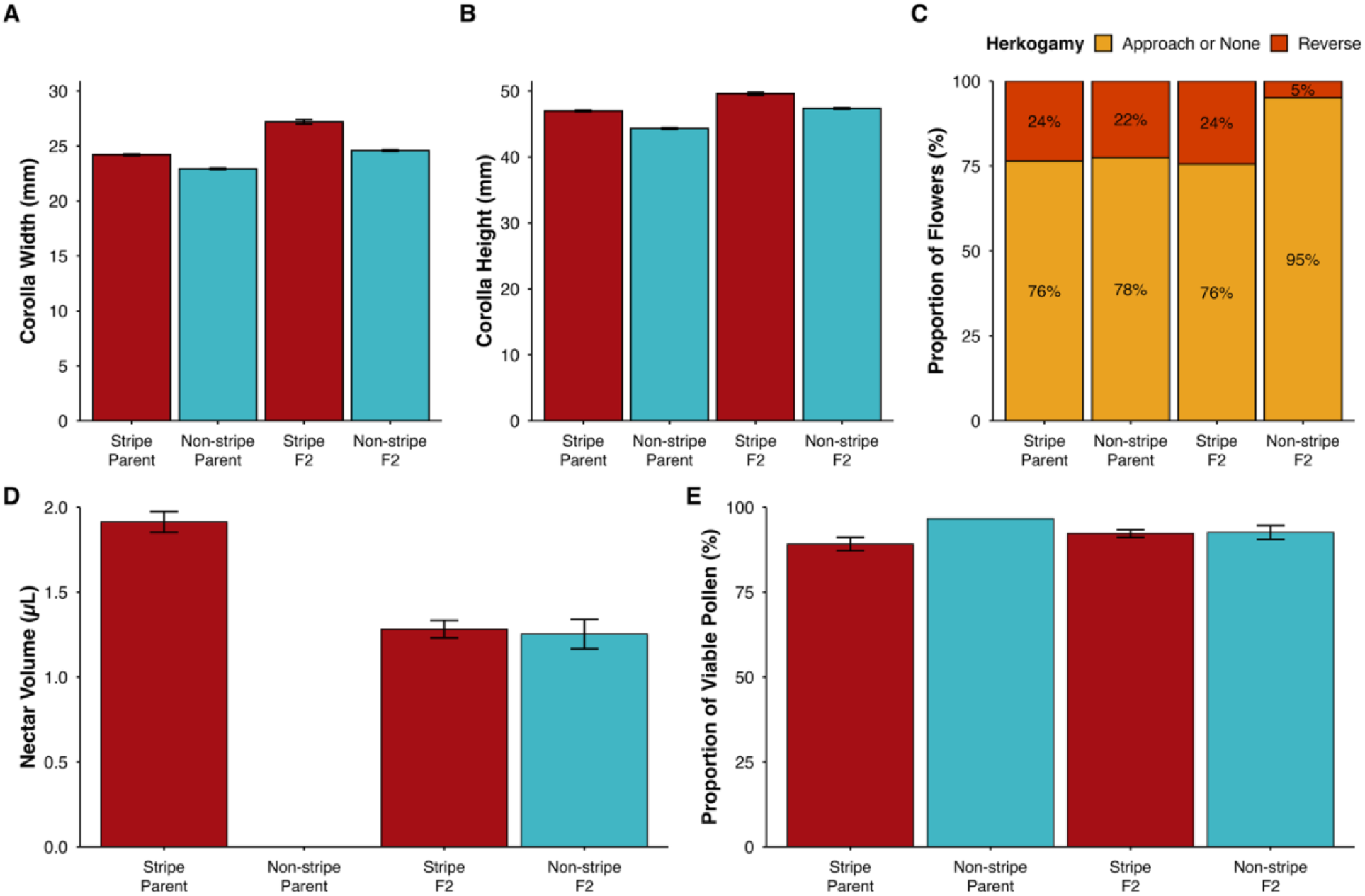
Floral traits of parent and F_2_ plants with and without foliar striping; error bars represent +/- 1 SE from the mean. (**A**) Corolla width, (**B**) Corolla height, and (**C**) Binary herkogamy (**D**) Nectar volume and (**E**) Pollen viability across groups.

**Table 1.** Results of analysis of variance demonstrating the effects of phenotype, genotype, and treatment (where appropriate) and their interaction on crimson monkeyflower traits. Any interactions not included here are due to their exclusion from the best fit models in (A) the field experiment and (B) the greenhouse experiment. Factors with significant effects are in bold type. Test-statistic is the F-statistic unless otherwise indicated.

| (A) |  | FIELD EXPERIMENT |  |  |  |
| --- | --- | --- | --- | --- | --- |
| Traits |  | Fixed Effects | DF<br>(numerator,<br>denominator) | Test-statistic | p-value |
| Floral | Corolla Width (mm) | **Phenotype** | 1, 471.5 | 110.09 | **< 0.001** |
|  |  | **Genotype** | 1, 441.34 | 171.44 | **< 0.001** |
|  |  | **P\*G Interaction** | 1, 471.51 | 10.76 | **< 0.01** |
|  | Corolla Height (mm) | **Phenotype** | 1, 470.9 | 67.43 | **< 0.001** |
|  |  | **Genotype** | 1, 440.9 | 100.19 | **< 0.001** |
|  |  | P*G Interaction | 1, 470.9 | 1.42 | 0.234 |
| | Herkogamy -<br>Reverse vs. Approach | **Phenotype** | 1 | $\chi^2 = 14.965$ | **< 0.001** |
| | | **Genotype** | 1 | $\chi^2 = 11.236$ | **< 0.001** |
| | | **P\*G Interaction** | 1 | $\chi^2 = 12.855$ | **< 0.001** |
| Fitness | Seed Set Presence/Absence | **Phenotype** | 1 | $\chi^2 = 8.912$ | **< 0.01** |
| | | Genotype | 1 | $\chi^2 = 2.102$ | 0.147 |
| | | **Treatment** | 1 | $\chi^2 = 59.628$ | **< 0.001** |
|  | Seed Number (of calyxes with seeds) | Phenotype | 1, 175.29 | 16.07 | < 0.001 |
|  |  | Genotype | 1, 170.64 | 4.66 | < 0.05 |
|  |  | Treatment | 1, 398.27 | 5.29 | < 0.05 |
|  |  | P*G*T Interaction | 1, 403.05 | 2.12 | 0.146 |
|  | Average Seed Weight per Individual Seed | Phenotype | 1, 120.10 | 4.91 | < 0.05 |
|  |  | Genotype | 1, 116.56 | 0.03 | 0.870 |
|  |  | Treatment | 1, 278.05 | 11.86 | < 0.001 |
|  | Average Seed Weight per Calyx | Phenotype | 1, 132.84 | 11.52 | < 0.001 |
|  |  | Genotype | 1, 129.56 | 0.80 | 0.373 |
|  |  | Treatment | 1, 291.94 | 23.48 | < 0.001 |
| PCA | PC1 | Phenotype | 2, 154 | t = -7.44 | < 0.001 |
|  |  | Genotype | 2, 154 | t = 5.78 | < 0.001 |
|  | PC2 | Phenotype | 2, 154 | t = -0.24 | 0.81 |
|  |  | Genotype | 2, 154 | t = 1.43 | 0.16 |
|  | PC3 | Phenotype | 2, 154 | t = 3.19 | < 0.01 |
|  |  | Genotype | 2, 154 | t = -7.77 | < 0.001 |
| (B) | GREENHOUSE EXPERIMENT |  |  |  |  |
| Traits |  | Fixed Effects | DF<br>(numerator,<br>denominator) | Test-statistic | p-value |
| Floral | Nectar Volume (uL) | Phenotype | 1, 186 | 0.0014 | 0.971 |
|  |  | Genotype | 1, 186 | 17.331 | < 0.001 |
| | Pollen Viability (%) | Phenotype | 1 | $\chi^2 = 34.169$ | < 0.001 |
| | | Genotype | 1 | $\chi^2 = 2.498$ | 0.114 |
| | | P*G Interaction | 1 | $\chi^2 = 14.862$ | < 0.001 |

**Table 2.**
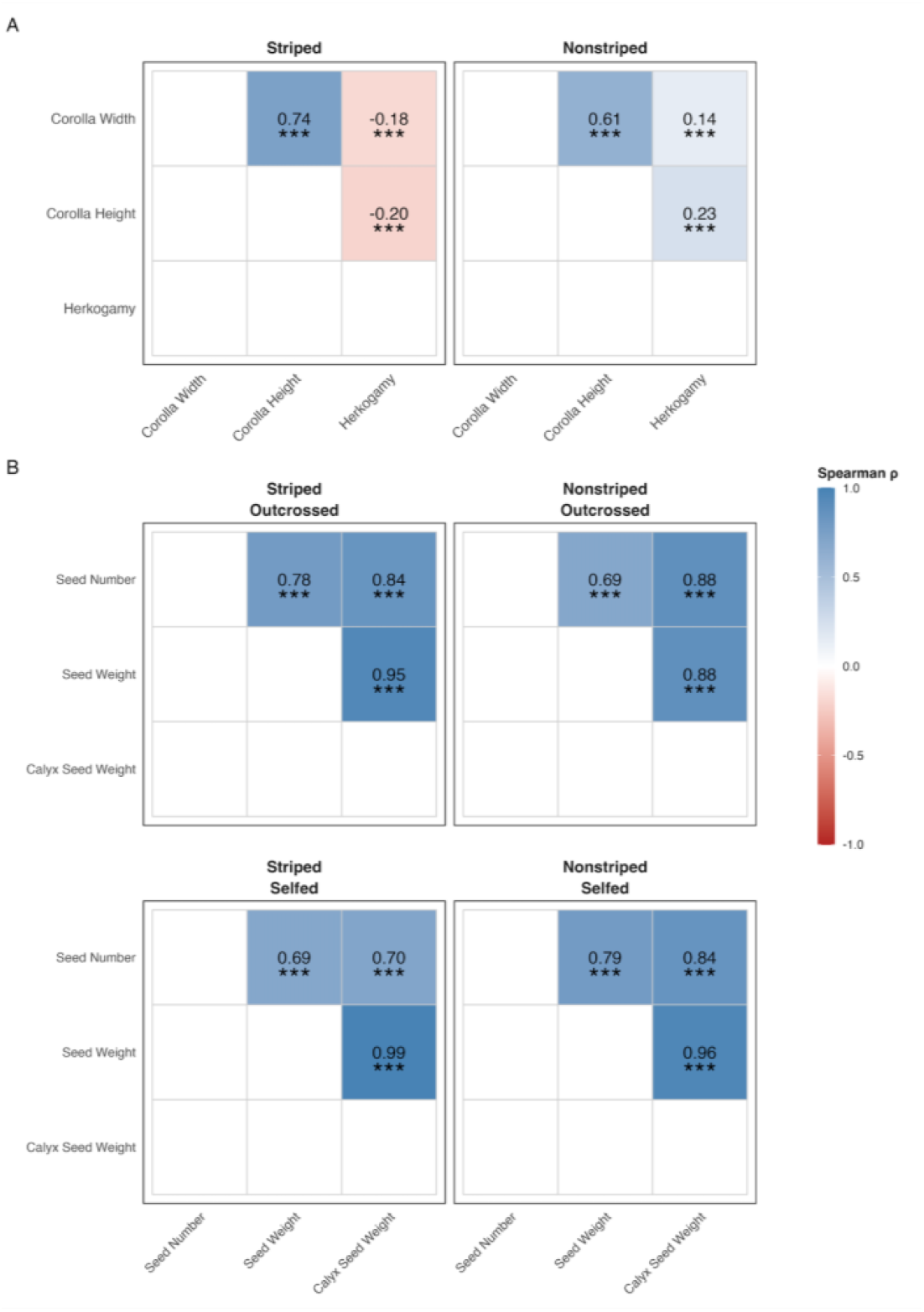
Full Spearman rank correlation matrices for field experiment data showing pairwise associations among (A) floral and (B) reproductive traits across foliar phenotypes and mating contexts, where experimentally appropriate. Correlations are displayed for all trait combinations regardless of statistical significance. Color indicates magnitude of Spearman’s ρ; asterisks denote FDR-adjusted significance (All correlations were significant at p < 0.001).

Corolla height was significantly related to both foliar striping phenotype and plant genotype, with striped plants producing corollas 1.05x taller than non-striped plants and parent plants producing corollas 0.94x the corolla height of F_2_ plants. For both corolla width and height, the greenhouse dataset showed largely similar patterns to the field experiment (Supplement Table 2; Supplement Figure 1).

### Herkogamy (Reverse vs Approach)

Across phenotypes, reverse herkogamy was generally rare; there was no difference in this trend between plants with presence or absence of foliar striping (Table 1A, 2A; Figure 2C), nor between parental and F_2_ genotypes. There was a significant phenotype x genotype interaction that was driven by F_2_ plants without foliar striping. While all parent plants and the F_2_ plants with foliar striping showed similarly low frequencies of reverse herkogamy (15-17%), F_2_ plants without foliar striping almost never expressed reverse herkogamy (2.5%), exhibiting 7-8x lower odds of reverse herkogamy than all other groups.

### Nectar Volume and Pollen Viability

As described in the Methods section, these traits were measured only in a subset of our greenhouse experiment plants. Under greenhouse conditions, the foliar striping phenotype was not related to nectar volume (Table 1B; Figure 2D) as foliar stripe and non-striped plants produced near identical volumes (< 1% difference). Genotype did significantly affect nectar production, as parental flowers produced 1.5x more nectar than F_2_ family flowers across foliar striping phenotypes (Figure 2D).

Pollen viability was significantly affected by the foliar striping phenotype and the phenotype by genotype interaction (Table 1B). Foliar striped plants had on average ∼3% lower pollen viability compared to non-striped plants (Figure 2E). The significant interaction between phenotype and genotype reflects the more pronounced difference in parent plants, relative to F_2_ plants, in pollen viability between plants with and without foliar striping. Striped parent plants had less than half the odds of viable pollen compared to the single non-striped parent plant assessed. Striped F_2_ plants had 15% lower odds of viable pollen than non-striped plants F_2_ plants. These patterns of pollen viability suggest that foliar striping is associated with lower male reproductive quality, independent of pollinator mediated processes, and that such a difference may reflect genetic or developmental tradeoffs.

**(Q2) (a) Is foliar striping associated with increased rates of pollination and/or pollination success, if so, does this (b) result in higher relative reproductive fitness in striped plants?**

### Seed set presence/absence

In our field experiment, individual flowers were given the opportunity to only self-fertilize via bagging, or to only outcross via external pollinators due to emasculation. Seed presence thus necessarily indicates successful self-fertilization or outcrossing. Seed presence was significantly influenced by both the foliar striping phenotype and mating system treatment (Table 1A). Plants with foliar striping were more than 1.35 times as likely to produce seed than non-striped plants, overall (Figure 3A). Obligately outcrossed flowers were more over 1.6 times more likely to set seed than obligately selfed flowers across phenotypes and genotypes (Figure 3A). There was not a significant phenotype x treatment interaction, indicating that plants with and without foliar striping respond similarly to the mating system treatments in terms of seed set success. Overall, these results demonstrate that foliar striping and obligate outcrossing are each associated with higher reproductive likelihood.

**Figure 3.**
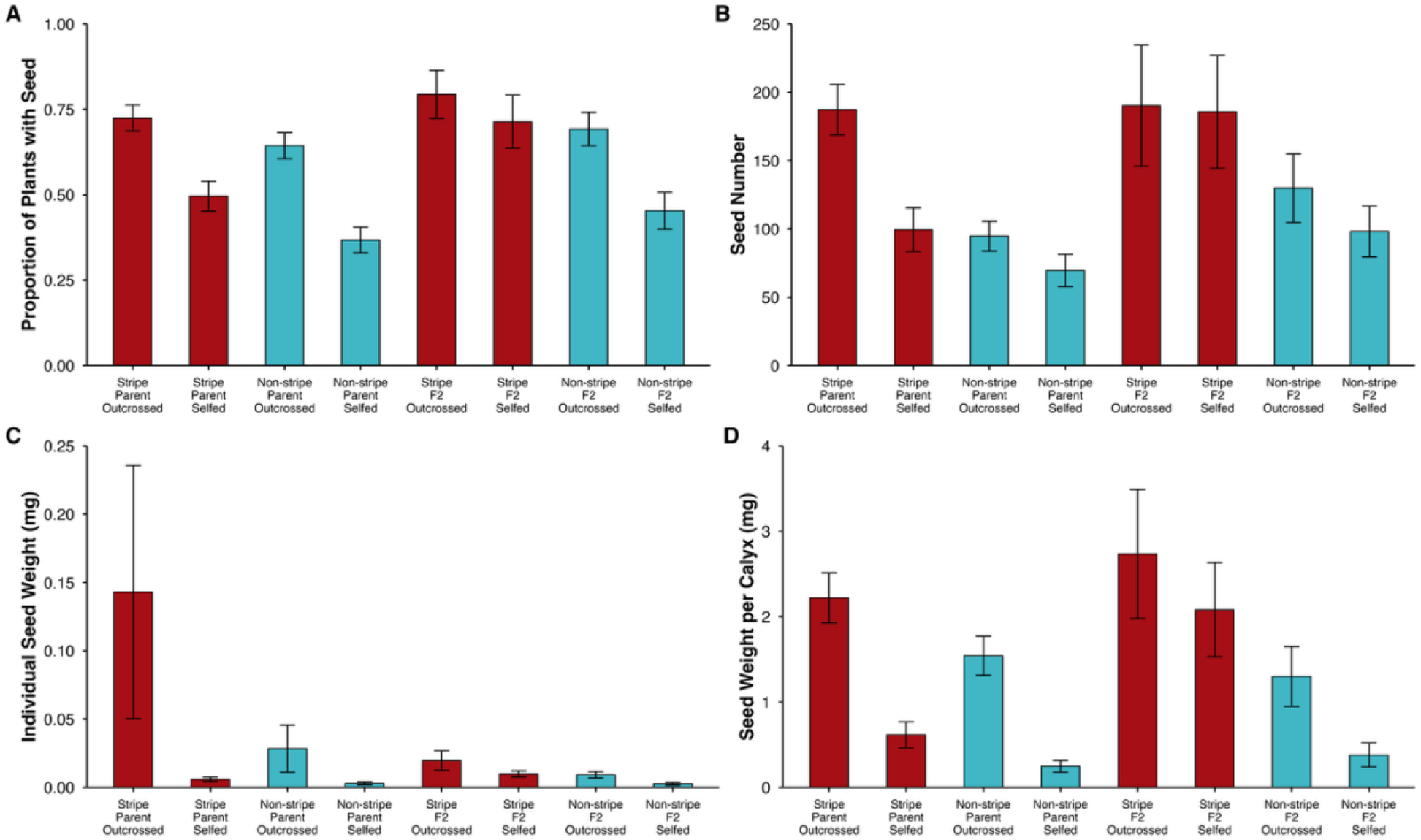
Seed set traits of plants with and without foliar striping, considering genotype and mating system treatment. All data is from the field experiment plants. Error bars represent +/- 1 SE from the mean. (**A**) Proportion of plants that produced seed, (**B**) Seed Number for those plants that did produce at least one seed, (**C**) Average seed weight (mg), (**D**) Total seed weight for all seeds in a calyx (mg).

**Figure 4.**
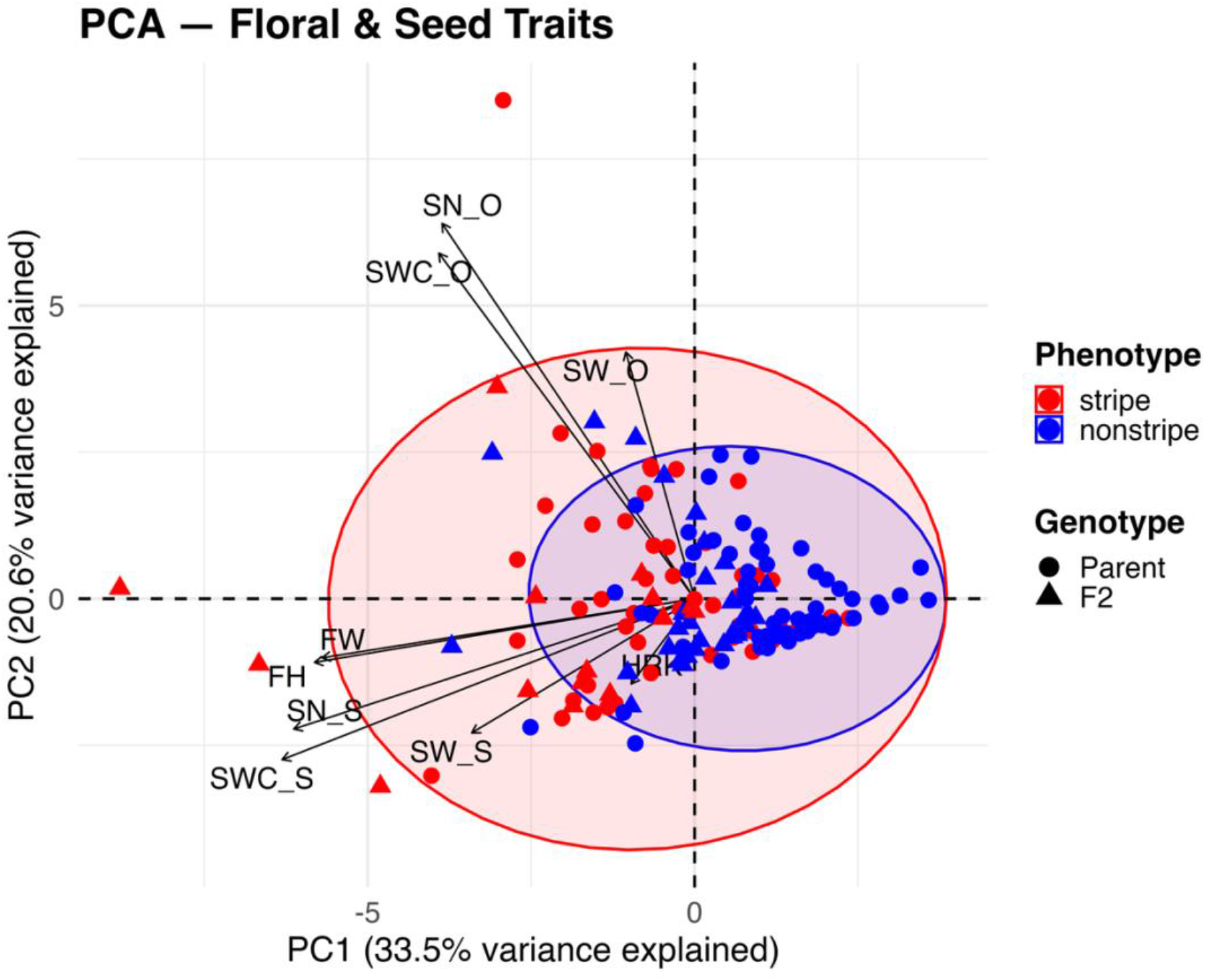
Combined floral and seed trait PCA from fieldwork data. Phenotype and Genotype are visually represented, though not included as part of the PC axes. Vector abbreviations are: (FW) flower width, (FH) flower height, (HRK) herkogamy, (SN_O) seed number-outcrossed, (SN_S) seed number-selfed, (SW_O) seed weight-outcrossed, (SW_S) seed weight-selfed, (SWC_O) seed weight calyx-outcrossed, (SWC_S) seed weight calyx-selfed.

### Seed Number

In addition to evaluating whether flowers produced seed in our field experiment, we examined seed number for flowers that successfully set seed. Unlike greenhouse conditions, where there was not pollen limitation, variation in seed number in the field reflects reproductive output per animal pollination event and provides information about pollination effectiveness and fertilization success. In the field, average seed number per calyx was 168 in plants with foliar stripes; 1.71 times higher than in flowers from non-stripe plants (Table 1A, 2B; Figure 3B). Genotype also affected seed number with flowers from parent plants producing 25% fewer seeds than flowers from F_2_ plants, regardless of foliar striping phenotype or mating system treatment. Mating system treatment significantly influenced seed production. Outcrossed flowers produced 1.31x more seeds than selfed flowers across phenotypes and genotypes. No significant interaction between foliar striping phenotype and mating system treatment was detected, indicating that regardless of foliar striping phenotype, plants experienced similar proportional increases in seed number under outcrossing relative to selfing.

Seed number from plants in the greenhouse experiment was significantly influenced only by the foliar striping phenotype (Supplemental Table 1; Supplemental Figure 2). Foliar striped plants produced 637 seeds per calyx on average; 1.33x more than in non-striped plants. Though outcrossed plants produced more seeds than selfed plants in the greenhouse, this trend was not statistically significant.

Across the field and greenhouse experiments, foliar striping was consistently associated with an increase in seed number per calyx, regardless of environment. Outcrossing via animal pollination was more effective than self-fertilization in the field environment. However, outcrossing via animal pollination did not produce nearly as many seeds per pollination event as when pollen was not limiting in the greenhouse hand pollinations.

### Seed Mass

#### Average Seed Mass per Individual Seed

Average individual seed mass serves as a proxy for per-offspring maternal investment and is commonly associated with early seedling performance (Roach and Wulff 1987). In our field experiment, average individual seed mass in plants with foliar striping was 0.017 mg; significantly greater than in flowers from non-stripe plants (Table 1A, 2B; Figure 3C). Across all genotypes and treatments, seeds produced by striped plants were 1.50x heavier than those produced by non-stripe plants. Genotype did not have a strong independent effect on average individual seed mass after accounting for phenotype and mating system treatment. Mating system treatment significantly affected individual seed mass. Obligately outcrossed flowers produced seeds that were, on average, 1.72x heavier than seeds produced by obligately self-fertilized flowers across phenotypes and genotypes. No significant interactions among foliar striping phenotype, genotype, or mating system treatment were detected for average individual seed mass, indicating that these factors act independently to influence the response of individual seed mass.

In our greenhouse experiment, average individual seed mass was significantly affected by phenotype, genotype, mating system, and the phenotype by genotype interaction (Supplemental Table 2; Supplemental Figure 2B). Plants with foliar striping produced seeds that were 0.014 mg, on average. In a pattern opposite to that in the field experiment, foliar striped plants produced 5.3% lighter seeds than non-striped plants. Parent plants produced seeds that were 1.14x heavier than seeds of F_2_ plants. As in the field experiment, outcrossed flowers produced heavier seeds than self-fertilization, but with a smaller difference: in the greenhouse seeds from outcrossed flowers were 1.26x heavier than those from selfed flowers. The phenotype by genotype interaction was driven largely by non-striped parent plants; these produced seeds that were ∼1.2x heavier than all other phenotype by genotype pairings.

Our individual seed mass results indicate that both foliar striping and outcrossing are independently associated with increased per-offspring investment in the field, with the seeds with the greatest mass produced by outcrossed flowers from plants with foliar striping.

#### Seed Mass per Calyx

Total seed mass per calyx (reproductive clutch) represents cumulative reproductive investment and integrates variation in both seed number and individual seed mass. In the field experiment, seed mass per calyx was significantly higher in flowers from plants with foliar striping than in flowers from non-stripe plants (Table 1A, 2B; Figure 3D). Reproductive clutches from striped plants weighed 2.64 mg on average, which was 1.75x more than those from non-stripe plants across genotypes and treatments. Parents and F_2_ did not differ in reproductive clutch mass. Mating system treatment had a strong effect on total reproductive investment. Outcrossed flowers produced calyces with about twice the total seed mass of selfed flowers across phenotypes and genotypes. No significant interactions among foliar striping phenotype, genotype, or mating system treatment were detected, indicating that reproductive clutch results are independently affected by each of these effects. Greenhouse results showed similar directional effects of phenotype and mating system but reduced phenotypic divergence (Supplemental Figure 2C).

### Trait Correlations

#### Correlation Matrix

Corolla width and corolla height were positively correlated across phenotypes (striped: ρ = 0.74; non-striped: ρ = 0.61), indicating that these floral size traits covary together (Table 2A). However, relationships between corolla size and herkogamy varied in direction across foliar phenotypes. In foliar striped plants, increases in corolla size were associated with shifts towards more negative herkogamy values (i.e., anthers further below the stigma), while increases in corolla size in non-striped plants were associated with shifts towards more positive herkogamy values (i.e., anthers above the stigma). While significant, these relationships had a relatively small effect size (|ρ| ≤ 0.23).

Reproductive traits were highly and consistently positively correlated across phenotypes and mating treatments in the field (Table 2B). Plants that produced more seed tended to produce seeds that are heavier, which contributed to heavier overall reproductive mass per calyx. This pattern was consistent across both striped and non-striped plants and under both mating system contexts (ρ ≈ 0.69-0.99).

#### Principal Component Analysis

The PCA of field experiment traits included corolla width, corolla height, herkogamy, selfed and outcrossed seed number, selfed and outcrossed average seed mass, and selfed and outcrossed calyx seed mass. The first two principal components explained 54.1% of total variation (PC1 = 33.5%, PC2 = 20.6%; Figure 5).

PC1 represented the dominant axis of trait covariation between floral size traits (corolla width and height) and selfed reproductive traits (selfed seed number, selfed average seed mass, selfed calyx seed mass). These traits strongly loaded together and in the negative direction, indicating that plants with larger corolla sizes tended to produce more seeds and greater seed mass in the self-fertilization treatment than plants with smaller corolla sizes.

PC2 captured variation primarily in outcrossed reproductive traits. Outcrossed seed number and outcrossed calyx seed mass loaded strongly and positively, with outcrossed average seed mass loading moderately. Floral morphology traits contributed less to PC2. This pattern indicates that variation in reproductive fitness via outcrossing is at least partially independent of floral size.

Individuals clustered broadly by phenotype along PC1, with striped and non-striped plants partially separated but overlapping. Variation along PC2 did not strongly separate by phenotypes, indicating that differences in outcrossed reproductive output occurred within both phenotype groups.

Herkogamy contributed weakly to both PC axes, which was consistent with the weak pairwise correlations between corolla size and herkogamy. Overall, the PCA results mirrored the correlation matrix patterns for floral and reproductive traits respectively.

## DISCUSSION

Through field and greenhouse experiments, we show that foliar anthocyanin striping in crimson monkeyflower is genetically correlated with floral traits associated with pollinator recruitment and reproductive success. Foliar striped plants generally produced larger corollas, more seeds and greater total seed mass per calyx relative to plants without foliar stripes, particularly under field conditions when outcrossed by pollinators. These patterns were more pronounced between stiped and non-striped F_2_ individuals than between parent lines, rather than being decoupled via independent assortment. In contrast, nectar volume, herkogamy, pollen viability, and individual seed mass did not co-vary with foliar striping but instead depended on genetic background and/or experimental environment (field or greenhouse). Our results demonstrate that foliar striping is part of a partially integrated foliar-floral phenotype, with some, but not all, floral traits showing clear phenotypic and genetic associations with foliar striping.

### Foliar-floral trait integration

Foliar striping was genetically associated with some, but not all, components of the floral phenotypes that may be relevant to pollination; these results are consistent with foliar striping being part of a pollination syndrome. Pollination syndromes are commonly described as combinations of floral traits, including flower size, shape, color, scent, and nectar rewards, that may work together to recruit particular pollinators (Fenster et al. 2004; Ollerton et al. 2009; Dellinger 2020). In both the parent lines and in the F_2_ individuals, foliar striped plants generally produced larger corollas, connecting the vegetative phenotype with floral display size. This association is notable because the null expectation, with empirical evidence from other systems, is that floral and vegetative traits should vary independently. For example, a synthesis study comparing trait correlations across 36 flowering plant species, found that correlations between floral and vegetative traits were weaker between groups than they were within groups (Connor et al. 2014).

### Pollinators

Hummingbirds were the primary floral visitors observed in our field arrays (E.R.L., personal observation). They moved among flowering plants, contacted the reproductive structures, and accumulated visible pollen on their foreheads. Rufous and Broad-tailed Hummingbirds were observed to defend the arrays from other visitors, including Anna’s Hummingbirds and occasionally White-lined Sphinx Moths. Territorial hummingbirds tend to move pollen within populations more frequently than between populations, as they focus on defending small, high-resource patches (Hadley et al. 2017). An early demonstration of this was in *Heliconia*, where pollen dispersal was more restricted where flowers occurred in dense patches defended by territorial hummingbirds (Linhart 1973). In natural populations of crimson monkeyflower, pollinator territoriality could result in increases in the level of inbreeding through near-neighbor transfer of pollen, or through self-pollination if multiple flowers on the same plant are visited in succession (Wessinger 2021).

### Foliar striping and pollinator behavior

We hypothesize that the foliar stripe could be contributing to whole-plant pollinator display by changing how the plant appears to visitors, as it forms a “bullseye” pattern around the corolla against the plant’s green leaves and the surrounding habitat. Hummingbirds can distinguish color combinations that include wavelengths outside of human vision; prior work in other systems supports the likelihood that they can distinguish between the foliar striped and non-striped phenotypes visually. For example, wild Broad-tailed Hummingbirds were trained to choose between paired LED-lit feeders, one containing sugar water and the other plain water, while repeatedly changing feeder position (Stoddard et al. 2020). The hummingbirds distinguished purple and several color combinations containing ultraviolet light, showing that they can learn visual differences that humans perceive differently or cannot see at all.

The extent to which the foliar stripe visually stands out likely depends on its contrast with the surrounding background, as well as the pollinator type, as the same signal can appear differently to different pollinators (Rodriguez-Girones and Santamaria 2004; Shrestha et al. 2019). In a community ecology study of bee- and hummingbird-pollinated plants, floral reflectance and color were modeled against the vegetative background as the respective pollinators would see them (de Camargo et al. 2019). Specific bee- and hummingbird-pollinated flowers were visually similar to the hummingbirds, but the hummingbird-pollinated flowers were less conspicuous to the bee pollinators. Bees have been shown to take longer to detect red flowers than others; this is hypothesized to be a result of red and green color receptors in bees having overlap in detection sensitivity peaks (Chittka and Waser 1997; Briscoe and Chittka 2001). It is possible that the foliar striping in our study species makes flowers less visible to bees than flowers with a purely green foliar background, if it reduces the visual contrast between the red flowers and green foliage (Spaethe et al. 2001; Rodriguez-Girones and Santamaria 2004). While we did not observe bee visitors to our study species in the field, small numbers of them have been observed to visit a sister species, *Mimulus cardinalis* and its hybrids (Schemske and Bradshaw 1999; Peng et al. 2024). Reduced attraction to bees relative to hummingbirds could increase the changes of successful outcrossing, according to the bee-avoidance hypothesis (Raven 1972; Oliveira et al. 2026). Reflectance spectroscopy of our experimental plants showed no UV reflectance from either the green leaf tissue or the striped tissue, and UV photography showed no distinct UV patterning between the two tissue regions (Supplementary Figure 4A, B). Therefore, any visual effect of the foliar stripe on pollinators is likely to come from the color spectrum or contrast at visible wavelengths rather than from a separate UV pattern.

While relatively rare, studies in other systems show that foliar traits can influence pollinator behavior. For example, in the wetland perennial Asian lizard’s tail (*Saururus chinensis)*, leaves surrounding the flowers turn white during flowering. When these white leaves were experimentally removed or covered, plants received fewer pollinator visits and produced fewer fruits and seeds (Song et al. 2018). Similarly, dove tree (*Davidia involucrata)* produces large white bracts around its flowers. When these white bracts were manually replaced with green ones or removed, bee attraction decreased compared to plants displaying natural or artificial white bracts (Sun et al. 2008). While both of these examples involve white foliar tissue, other work has shown similar examples with anthocyanin-pigmented bracts (Bergamo et al. 2019).

### Female reproductive traits

Corolla size may affect both whether a hummingbird visits a plant and how well pollen is transferred by visiting pollinators. For example, in an experimental assessment of pollen transfer in scarlet gilia (*Ipomopsis aggregata)*, hummingbirds inserted their bills more deeply into wider flowers and removed a greater proportion of the available pollen (Campbell et al. 1996). The larger corollas of striped, crimson monkeyflowers may therefore contribute to pollinator attraction and/or pollen transfer alongside the foliar stripe. Our reproductive output results in the field experiment are compatible with either/both mechanisms, showing greater per calyx seed production via outcrossing in plants with foliar stripes and larger corollas.

In our field experiment, flowers in the outcrossing treatment were emasculated, so they could produce seeds only after pollen was delivered from another plant by a pollinator. The greater per-flower reproductive output of foliar striped plants, compared to non-striped plants, provides indirect evidence of pollinator preference for, and/or more effective pollination of, the integrated striped phenotype. This phenomenon could have multiple possible mechanisms. Prior pollen-supplementation experiments have found that receiving an increased pollen load increases fruit or seed production, demonstrating that under natural conditions the pollen received may limit reproduction through its quantity or quality (Ashman et al. 2004). Possible mechanisms underlying the outcrossed seed production differences in our experiment include the delivery of differential pollen load or viability from prior-visited flowers, differences in pollinator visitation rates, and/or effectiveness of pollen deposition (Opedal et al. 2023).

However, pollinator-mediated pollen delivery likely does not fully explain the greater per-calyx reproductive output of outcrossed foliar striped plants. Flowers that were obligately self-fertilizing were less likely to produce seeds than outcrossed flowers, but among those that did produce seeds, per-calyx selfed reproductive output was also higher in plants with foliar striping and (on average) larger corollas. It is possible that larger flowers have greater seed production capacity. Prior work in white campion (*Silene latifolia*) showed a positive genetic correlation with corolla size and number of ovules, and a negative genetic correlation between corolla size and per-plant flower number (Delph et al. 2004). Given that we did not assess the number of flowers produced per plant, we note that our per-flower reproductive output results do not necessarily translate to whole plant reproductive fitness.

Support for the hypothesis that flowers from plants with foliar striping and larger corollas have greater seed production capacity, comes from our greenhouse experiment. Greenhouse flowers were hand-pollinated in a standardized way that maximized the pollen load for each flower. In this way, our greenhouse results represent possible per flower reproductive capacity for our experimental plants when self-fertilized or outcrossed. Per-calyx seed production was greater in the greenhouse, relative to the field. Despite the difference in scale, greenhouse plants showed the same pattern of plants with foliar stripes producing more seeds than non-striped plants. In a small subset of F_2_ plants in which both traits were measured, corolla width was positively associated with flower count (F1,24 = 10.8; p-value =0.003).

We did not find trade-offs between seed number and seed mass per calyx in the field experiment, in contrast to classic theory that predicts that with a finite amount of resources, an individual will produce fewer large seeds or more, smaller seeds (Smith and Fretwell 1974; Fagundes et al. 2025). For example, in a study of floral size and seed production in the perennial herb, *Paeonia broteroi,* larger corollas produced more, but lighter, seeds (Sánchez-Lafuente and Parra 2009). Adherence to this prediction depends on resource availability, amongst other factors. In our study, plants were placed in the field environment in their greenhouse pots, potentially reducing nutrient-related stressors that would be present in the natural soil environment.

### Male reproductive traits

In contrast to their increased display size and higher seed production, aspects of female reproductive fitness, flowers from foliar striped plants had slightly lower pollen viability overall. This pattern was largely due to differences between the parent plants, rather than between F_2_ individuals with and without foliar striping, indicating that this trait may be assorting independently with foliar striping. Producing functional pollen is only the one aspect of male reproductive fitness. The rate at which pollen leaves the anther, is carried to and reaches the stigma of another flower are relevant here (Opedal et al. 2023) and may be influenced by floral display and/or foliar striping. Thus, our results identify a difference in pollen quality rather than necessarily a difference in total male fitness.

### Mating system

The ability to produce seeds through both self-fertilization and outcrossing can provide reproductive assurance when compatible pollen or pollinator visits are limited, while outcrossing may reduce the genetic costs associated with inbreeding (Herlihy and Eckert 2002; Kalisz et al. 2004; Goodwillie et al. 2005). While many flowers in our field experiment did successfully self-fertilize, per-calyx seed number and mass were higher from outcrossed flowers. This trend was similar in the greenhouse experiment (although significant only for per-calyx seed mass) indicating that the relative decrease in seed production across mating systems is not simply a result of less effective pollen delivery in naturally self-fertilizing flowers. The approach herkogamy observed in the great majority of flowers in our experiments supports the idea that delayed selfing is a method of reproductive assurance in these crimson monkeyflower plants, similarly to many other species (Busch and Delph 2012; Goodwillie and Weber 2018).

### Future directions

Our study demonstrates that foliar striping and genetically correlated traits are associated with increased per-calyx reproductive output in crimson monkeyflower under both natural and artificial pollination. The full evolutionary implications of this phenomenon could be elucidated by future work examining whole plant reproductive fitness via field experiments. In addition, future work focused on pollinator behavior and associated plant cues could clarify the effects of individual traits on pollinator visitation patterns. Measuring reflectance across the full spectrum and modeling the foliar stripe against the leaves and surrounding habitat would show how strong the contrast is from the perspective of different pollinators.

## ACKNOWLEDGMENTS

Support for this research was provided by NIH R01GM131055, NSF-REU Site 2348925), and NAU Faculty Research Grants program. Thanks to Yao-Wu Yuan and Amy LaFountain for their support and discussions surrounding this work. Permit Number for fieldwork and seed collections (collections permit: FS-2400-008 (permit number RO2021.21); field site permit: FS-2700-4). Thank you to the Forest Service permitting office for their assistance, including Deirdre Marx, Janie Agyago, and Tobias J. Hutchens. Thank you to Kelsey Byers for running preliminary experiments on the reflectance spectroscopy and UV photography. Lara Schmidt in CAWL and the NAU Greenhouse, specifically Adair Patterson, provided resources, support, and expertise. Thank you to ERL’s PhD committee (Amy Whipple, Brad Butterfield, and Rich Hofstetter) for their feedback on this research. Rich Hofstetter first noted the bullseye-like pattern of foliar striping around the flower. Holeski Lab members provided edits and advice. The following undergraduate researchers contributed to this research (in chronological order): Jess Combs (REU), Eva Canby, Kellee Englert, Sada DeWitt, Joseph Gastelum (REU), and Madison Hughes. The following volunteers assisted with fieldwork and field observations: Audrey Harvey, Matthew Weiss, Dr. Greg Cruz, Shelley LaPlante, Shane LaPlante, and Kari LaPlante. A special thanks to Dean Lehman for the use of his private property entrance and parking to conveniently access field sites for 6-months.

ChatGPT (OpenAI, ChatGPT web application; GPT-5.2 Instant, GPT-5.3 Instant, GPT-5.5 Instant, GPT-5.5 Thinking, and GPT-5.6 Sol; January-August 2026) was used interactively to assist with brainstorming manuscript organization and developing and troubleshooting R code. Prompts included author-written manuscript passages, descriptions of the study design and analytical objectives, R scripts, and associated error messages. Model responses were treated as suggestions and were reviewed and revised by the authors. The authors independently verified all statistical methods, code, numerical results, and final manuscript text and take full responsibility for the submitted work.

## REFERENCES

Ahmed, N. U., Park, J.-I., Jung, H.-J., Yang, T.-J., Hur, Y., & Nou, I.-S. (2014). Characterization of dihydroflavonol 4-reductase (DFR) genes and their association with cold and freezing stress in *Brassica rapa*. Gene, 550(1), 46–55.

Altshuler, D. L. (2003). Flower color, hummingbird pollination, and habitat irradiance in four Neotropical forests. Biotropica, 35(3), 344–355.

Altshuler, D. L., & Dudley, R. (2002). The ecological and evolutionary interface of hummingbird flight physiology. Journal of Experimental Biology, 205(16), 2325–2336.

Altshuler, D. L., & Wylie, D. R. (2020). Hummingbird vision. Current Biology, 30(3), R103– R105.

Armbruster, W. S., Pélabon, C., Bolstad, G. H., & Hansen, T. F. (2014). Integrated phenotypes: understanding trait covariation in plants and animals. Philosophical Transactions of the Royal Society B: Biological Sciences, 369(1649), 20130245.

Armbruster, W. S., Corbet, S. A., Vey, A. J., Liu, S. J., & Huang, S. Q. (2014). In the right place at the right time: Parnassia resolves the herkogamy dilemma by accurate repositioning of stamens and stigmas. Annals of botany, 113(1), 97–103.

Ashman, T.-L., Knight, T. M., Steets, J. A., Amarasekare, P., Burd, M., Campbell, D. R., Dudash, M. R., Johnston, M. O., Mazer, S. J., Mitchell, R. J., Morgan, M. T., & Wilson, W. G. (2004). Pollen limitation of plant reproduction: Ecological and evolutionary causes and consequences. Ecology, 85(9), 2408–2421.

Barrett, S. C. H. (2003). Mating strategies in flowering plants: The outcrossing–selfing paradigm and beyond. Philosophical Transactions of the Royal Society of London. Series B: Biological Sciences, 358(1434), 991–1004.

Bates, D., Mächler, M., Bolker, B., & Walker, S. (2015). Fitting linear mixed-effects models using lme4. Journal of Statistical Software, 67(1), 1–48.

Benzing, D. H., & Friedman, W. E. (1981). Patterns of foliar pigmentation in Bromeliaceae and their adaptive significance. Selbyana, 5(3/4), 224–240.

Bergamo, P. J., Wolowski, M., Telles, F. J., De Brito, V. L. G., Varassin, I. G., & Sazima, M. (2019). Bracts and long-tube flowers of hummingbird-pollinated plants are conspicuous to hummingbirds but not to bees. Biological Journal of the Linnean Society, 126(3), 533–544.

Briscoe AD, Chittka L. (2001). The evolution of color vision in insects. Annu Rev Entomol, 46, 471–510.

Busch, J. W., & Delph, L. F. (2012). The relative importance of reproductive assurance and automatic selection as hypotheses for the evolution of self-fertilization. Annals of botany, 109(3), 553–562.

Campbell, D. R. (1989). Measurements of selection in a hermaphroditic plant: Variation in male and female pollination success. Evolution, 43(2), 318–334.

Campbell, D. R., Waser, N. M., & Price, M. V. (1996). Mechanisms of hummingbird-mediated selection for flower width in *Ipomopsis aggregata*. Ecology, 77(5), 1463–1472.

Carr, D. E., & Dudash, M. R. (1997). The effects of five generations of enforced selfing on potential male and female function in *Mimulus guttatus*. Evolution, 51(6), 1797–1807.

Chalker-Scott, L. (1999). Environmental significance of anthocyanins in plant stress responses. Photochemistry and Photobiology, 70(1), 1–9.

Chen, H., Berg, C. S., Samuli, M., Sotola, V. A., Sweigart, A. L., Yuan, Y.-W., & Fishman, L. (2025). The genetic architecture of floral trait divergence between hummingbird- and self-pollinated monkeyflower (*Mimulus*) species. New Phytologist, 245(5), 2255–2267.

Chittka, L., & Waser, N. M. (1997). Why red flowers are not invisible to bees. Israel Journal of Plant Sciences, 45(2-3), 169–183.

Damián, X., Ochoa-López, S., Gaxiola, A., Fornoni, J., Domínguez, C. A., & Boege, K. (2020). Natural selection acting on integrated phenotypes: Covariance among functional leaf traits increases plant fitness. New Phytologist, 225(1), 546–557.

de Camargo, M. G. G., Lunau, K., Batalha, M. A., Brings, S., de Brito, V. L. G., & Morellato, L. P. C. (2019). How flower colour signals allure bees and hummingbirds: A community-level test of the bee avoidance hypothesis. New Phytologist, 222(2), 1112–1122.

Dellinger, A. S. (2020). Pollination syndromes in the 21st century: Where do we stand and where may we go? New Phytologist, 228(4), 1193–1213.

Delph, L. F., Gehring, J. L., Frey, F. M., Arntz, A. M., & Levri, M. (2004). Genetic constraints on floral evolution in a sexually dimorphic plant revealed by artificial selection. Evolution, 58(9), 1936–1946.

Ding, B., Patterson, E. L., Holalu, S. V., Li, J., Johnson, G. A., Stanley, L. E., Greenlee, A. B., Peng, F., Bradshaw, H. D., Jr., Blinov, M. L., Blackman, B. K., & Yuan, Y.-W. (2020). Two MYB proteins in a self-organizing activator-inhibitor system produce spotted pigmentation patterns. Current Biology, 30(5), 802–814.

Faegri, K., & van der Pijl, L. (2013). Principles of pollination ecology. Springer.

Fagundes, M., Rodrigues, S. M., Cuevas-Reyes, P., Camarota, F., Jorge, A. C., & Figueiredo, L. H. A. (2025). Breaking the norm: Absence of seed size/seed number trade-off in Hymanaea stigonocarpa, a tree species from the Brazilian semi-arid. Plant Species Biology, 40(5), 472–482.

Fellows, T. K. (2015). Visual resolution of Anna’s hummingbirds (Calypte anna) in space and time [Master’s thesis, University of British Columbia]. UBC Library Open Collections.

Fenster, C. B. (1991). Selection on floral morphology by hummingbirds. Biotropica, 23(1), 98– 101.

Fenster, C. B., Armbruster, W. S., Wilson, P., Dudash, M. R., & Thomson, J. D. (2004). Pollination syndromes and floral specialization. Annual Review of Ecology, Evolution, and Systematics, 35(1), 375–403.

Goodwillie, C., & Weber, J. J. (2018). The best of both worlds? A review of delayed selfing in flowering plants. American Journal of Botany, 105(4), 641–655.

Goodwillie, C., Kalisz, S., & Eckert, C. G. (2005). The evolutionary enigma of mixed mating systems in plants: Occurrence, theoretical explanations, and empirical evidence. Annual Review of Ecology, Evolution, and Systematics, 36(1), 47–79.

Hadley, A. S., Frey, S. J., Robinson, W. D., & Betts, M. G. (2018). Forest fragmentation and loss reduce richness, availability, and specialization in tropical hummingbird communities. Biotropica, 50(1), 74–83.

Hartig, F. (2017). *DHARMa: Residual diagnostics for hierarchical (multi-level/mixed) regression models* [R package].

Herlihy, C. R., & Eckert, C. G. (2002). Genetic cost of reproductive assurance in a self-fertilizing plant. Nature, 416(6878), 320–323.

Hurly, T. A., & Healy, S. D. (1996). Memory for flowers in rufous hummingbirds: Location or local visual cues? Animal Behaviour, 51(5), 1149–1157.

iNaturalist. (n.d.). *Erythranthe verbenacea (crimson monkeyflower)*. Retrieved March 9, 2024, from [insert exact iNaturalist URL]

Jablonski, D. (2008). Species selection: Theory and data. Annual Review of Ecology, Evolution, and Systematics, 39(1), 501–524.

Kalisz, S., Vogler, D. W., & Hanley, K. M. (2004). Context-dependent autonomous self-fertilization yields reproductive assurance and mixed mating. Nature, 430(7002), 884– 887.

Kearns, C. A., & Inouye, D. W. (1993). Techniques for pollination biologists. University Press of Colorado.

Kessler, D., & Baldwin, I. T. (2015). How scent and nectar influence floral antagonists and mutualists. Functional Ecology, 29(3), 529–538.

Koski, M. H., Leonard, E., & Tharayil, N. (2024). Foliar flavonoids across an elevation gradient: Plasticity in response to UV, and links with floral pigmentation patterning. Environmental and Experimental Botany, 228, 106036.

LaFountain, A. M., & Yuan, Y.-W. (2021). Repressors of anthocyanin biosynthesis. New Phytologist, 231(3), 933–949.

LaFountain, A. M., McMahon, H. E., Reid, N. M., & Yuan, Y.-W. (2023). To stripe or not to stripe: The origin of a novel foliar pigmentation pattern in monkeyflowers (*Mimulus*). New Phytologist, 237(1), 310–322.

Lázaro, A., Seguí, J., & Santamaría, L. (2020). Continuous variation in herkogamy enhances the reproductive response of *Lonicera implexa* to spatial variation in pollinator assemblages. AoB PLANTS, 12(1), plz078.

Leimberger, K. G., Dalsgaard, B., Tobias, J. A., Wolf, C., & Betts, M. G. (2022). The evolution, ecology, and conservation of hummingbirds and their interactions with flowering plants. Biological Reviews, 97(3), 923–959.

Lenth, R. V. (2023). *emmeans: Estimated marginal means, aka least-squares means* (Version 1.8.5) [R package].

Lev-Yadun, S., & Gould, K. S. (2008). Role of anthocyanins in plant defence. In Anthocyanins: Biosynthesis, functions, and applications (pp. 22–28). Springer.

Lev-Yadun, S., Inbar, M., Izhaki, I., & Dafni, A. (2002). Colour patterns in vegetative parts of plants deserve more research attention. Trends in Plant Science, 7(2), 59–60.

Liang, M., Foster, C. E., & Yuan, Y.-W. (2022). Lost in translation: Molecular basis of reduced flower coloration in a self-pollinated monkeyflower (*Mimulus*) species. Science Advances, 8(37), eabo1113.

Linhart, Y. B. (1973). Ecological and behavioral determinants of pollen dispersal in hummingbird-pollinated *Heliconia*. The American Naturalist, 107(956), 511–523.

Lisney, T. J., Wylie, D. R., Kolominsky, J., & Iwaniuk, A. N. (2015). Eye morphology and retinal topography in hummingbirds (Trochilidae: Aves). Brain, Behavior and Evolution, 86(3–4), 176–190.

Lloyd, D. G., & Webb, C. J. (1986). The avoidance of interference between the presentation of pollen and stigmas in angiosperms. I. Dichogamy. New Zealand Journal of Botany, 24(1), 135–162.

Lowry, D. B., Sheng, C. C., Lasky, J. R., & Willis, J. H. (2012). Five anthocyanin polymorphisms are associated with an R2R3-MYB cluster in *Mimulus guttatus* (Phrymaceae). American Journal of Botany, 99(1), 82–91.

Meléndez-Ackerman, E., Campbell, D. R., & Waser, N. M. (1997). Hummingbird behavior and mechanisms of selection on flower color in *Ipomopsis*. Ecology, 78(8), 2532–2541.

Minnaar, C., Anderson, B., de Jager, M. L., & Karron, J. D. (2019). Plant–pollinator interactions along the pathway to paternity. Annals of Botany, 123(2), 225–245.

Nesom, G. L. (2014). Taxonomy of *Erythranthe* sect. Erythranthe (Phrymaceae). Phytoneuron, 2014(31), 1–41.

Oliveira, L. C., Brito, V. L. G., Lunau, K., Gerten, S., Oliveira, P. E. M., Melo, L. R. F., Telles, F. J., & Bergamo, P. J. (2026). Evolution of UV reflection in bee-and bird-pollinated flowers. Plant Biology, 28(1), 201–214.

Ollerton, J., Alarcón, R., Waser, N. M., Price, M. V., Watts, S., Cranmer, L., Hingston, A., Peter, C. I., & Rotenberry, J. (2009). A global test of the pollination syndrome hypothesis. Annals of botany, 103(9), 1471–1480.

Opedal, Ø. H., Pérez-Barrales, R., Brito, V. L. G., Muchhala, N., Capó, M., & Dellinger, A. S. (2023). Pollen as the link between floral phenotype and fitness. American Journal of Botany, 110(6), e16200.

Peng, F., Sun, X., van Vloten, C., Correll, J., Langdon, M., Ngochanthra, W., Johnson, K., & Amador Kane, S. (2024). Hybrid Mimulus flowers attract a new pollinator. New Phytologist, 242(3), 1324–1332.

Peng, F., Byers, K. J. R. P., & Bradshaw, H. D., Jr. (2017). Less is more: Independent loss-of-function *OCIMENE SYNTHASE* alleles parallel pollination syndrome diversification in monkeyflowers (*Mimulus*). American Journal of Botany, 104(7), 1055–1059.

Phillips, P. C., & Arnold, S. J. (1989). Visualizing multivariate selection. Evolution, 43(6), 1209–1222.

Pigliucci, M. (2003). Phenotypic integration: Studying the ecology and evolution of complex phenotypes. Ecology Letters, 6(3), 265–272.

R Core Team. (2023). R: A language and environment for statistical computing. R Foundation for Statistical Computing. https://www.R-project.org/

Raven, P. H. (1972). Why are bird-visited flowers predominantly red?. Evolution, 26(4), 674–674.

Roach, D. A., & Wulff, R. D. (1987). Maternal effects in plants. Annual Review of Ecology and Systematics, 18, 209–235.

Rodríguez-Gironés, M. A., & Santamaría, L. (2004). Why are so many bird flowers red?. PLoS biology, 2(10), e350.

Sánchez-Lafuente, A. M., & Parra, R. (2009). Implications of a long-term, pollinator-mediated selection on floral traits in a generalist herb. Annals of Botany, 104(4), 689–701.

Schemske, D. W., & Bradshaw, H. D., Jr. (1999). Pollinator preference and the evolution of floral traits in monkeyflowers (*Mimulus*). Proceedings of the National Academy of Sciences of the United States of America, 96(21), 11910–11915.

Shrestha, M., Dyer, A. G., Garcia, J. E., & Burd, M. (2019). Floral colour structure in two Australian herbaceous communities: it depends on who is looking. Annals of Botany, 124(2), 221–232.

Simpson, G. G. (1944). Tempo and mode in evolution. Columbia University Press.

Sinnott-Armstrong, M. A., Deanna, R., Pretz, C., Liu, S., Harris, J. C., Dunbar-Wallis, A., Smith, S. D., & Wheeler, L. C. (2022). How to approach the study of syndromes in macroevolution and ecology. Ecology and Evolution, 12(3), e8583.

Smith, C. C., & Fretwell, S. D. (1974). The optimal balance between size and number of offspring. The American Naturalist, 108(962), 499–506.

Song, B., Stöcklin, J., Armbruster, W. S., Gao, Y., Peng, D., & Sun, H. (2018). Reversible colour change in leaves enhances pollinator attraction and reproductive success in *Saururus chinensis* (Saururaceae). Annals of Botany, 121(4), 641–650.

Spaethe, J., Tautz, J., & Chittka, L. (2001). Visual constraints in foraging bumblebees: flower size and color affect search time and flight behavior. Proceedings of the National Academy of Sciences, 98(7), 3898–3903.

Stoddard, M. C., Eyster, H. N., Hogan, B. G., Morris, D. H., Soucy, E. R., & Inouye, D. W. (2020). Wild hummingbirds discriminate nonspectral colors. Proceedings of the National Academy of Sciences of the United States of America, 117(26), 15112–15122.

Sun, J. F., Gong, Y. B., Renner, S. S., & Huang, S. Q. (2008). Multifunctional bracts in the dove tree *Davidia involucrata* (Nyssaceae: Cornales): Rain protection and pollinator attraction. The American Naturalist, 171(1), 119–124.

Vickery, R. K., Jr. (2008). How does *Mimulus verbenaceus* (Phrymaceae) set seed in the absence of pollinators? Evolutionary Biology, 35(3), 199–207.

Vickery, R. K., Jr., & Vickery, P. K., Jr. (1992). Pollinator preferences for yellow, orange, and red flowers of *Mimulus verbenaceus* and *M. cardinalis*. The Great Basin Naturalist, 52(2), 145–148.

Walsh, B., & Blows, M. W. (2009). Abundant genetic variation + strong selection = multivariate genetic constraints: A geometric view of adaptation. Annual Review of Ecology, Evolution, and Systematics, 40(1), 41–59.

Webb, C. J., & Lloyd, D. G. (1986). The avoidance of interference between the presentation of pollen and stigmas in angiosperms. II. Herkogamy. New Zealand Journal of Botany, 24(1), 163–178.

Weiss, M., Faske, T. M., & Holeski, L. M. (2026). Climate Gradients and Habitat Discontinuity Structure Genetic Variation in a Spring-Specialist Plant. bioRxiv, 2026-05.

Wenzell, K. E., Neequaye, M., Paajanen, P., Hill, L., Brett, P., & Byers, K. J. R. P. (2025). Within-species floral evolution reveals convergence in adaptive walks during incipient pollinator shift. Nature Communications, 16(1), 1–20.

Wessinger, C. A. (2021). From pollen dispersal to plant diversification: genetic consequences of pollination mode. New Phytologist, 229(6), 3125–3132.

Wright, S. (1931). Evolution in Mendelian populations. Genetics, 16(2), 97–159.

Ye, Z. M., Jin, X. F., Yang, J., Wang, Q. F., & Yang, C. F. (2019). Accurate position exchange of stamen and stigma by movement in opposite directions resolves the herkogamy dilemma in a protandrous plant, *Ajuga decumbens* (Labiatae). AoB PLANTS, 11(5), plz052.

Young, T. P. (1982). Bird visitation, seed-set, and germination rates in two species of *Lobelia* on Mount Kenya. Ecology, 63(6), 1983–1986.

Yuan, Y.-W. (2019). Monkeyflowers (*Mimulus*): New model for plant developmental genetics and evo-devo. New Phytologist, 222(2), 694–700.

Yuan, Y.-W., Rebocho, A. B., Sagawa, J. M., Stanley, L. E., & Bradshaw, H. D., Jr. (2016). Competition between anthocyanin and flavonol biosynthesis produces spatial pattern variation of floral pigments between *Mimulus* species. Proceedings of the National Academy of Sciences of the United States of America, 113(9), 2448–2453.

Yuan, Y.-W., Sagawa, J. M., Frost, L., Vela, J. P., & Bradshaw, H. D., Jr. (2014). Transcriptional control of floral anthocyanin pigmentation in monkeyflowers (*Mimulus*). New Phytologist, 204(4), 1013–1027.

Yuan, Y.-W., Sagawa, J. M., Young, R. C., Christensen, B. J., & Bradshaw, H. D., Jr. (2013). Genetic dissection of a major anthocyanin QTL contributing to pollinator-mediated reproductive isolation between sister species of *Mimulus*. Genetics, 194(1), 255–263.

