## Supplemental for "Foliar striping and floral traits are correlated with higher reproductive output in crimson monkeyflowers"

**Supplemental Table 1.** (**A**) Sample sizes for floral and seed set traits used in the fieldwork experiment, considering phenotype and genotype. Fieldwork sample sizes are Corolla height: n = 1,809, Corolla width: n = 1,809, Herkogamy: n = 1,809, and seed set traits, seed number n=841 and average seed weight n=623. (**B**) Sample sizes for floral and seed set traits used in the greenhouse experiments, considering phenotype and genotype. Greenhouse sample sizes are Corolla height: n = 915, Corolla width: n = 915, Herkogamy: n = 915, Nectar volume: n = 189, Pollen viability: n = 153, and seed set traits n=164.

| (A) | **FIELD EXPERIMENT** | |
|---|---|---|
| | *Floral Traits* | |
| Foliar Phenotype | Corolla Width, Corolla Height, & Herkogamy | |
| | **Parent** | **$F_2$** |
| **Stripe** | 663 | 82 |
| **Non-stripe** | 655 | 409 |

| | *Fitness Traits* | | | |
|---|---|---|---|---|
| Foliar Phenotype | Seed Number | | Average Seed Weight (per individual seed and per calyx) | |
| | **Parent** | **$F_2$** | **Parent** | **$F_2$** |
| **Stripe** | 269 | 69 | 206 | 62 |
| **Non-stripe** | 326 | 177 | 133 | 222 |

| (B) | **GREENHOUSE EXPERIMENTS** | | | | | |
|---|---|---|---|---|---|---|
| | *Floral Traits* | | | | | |
| Foliar Phenotype | Corolla Width, Height, & Herkogamy | | Nectar Volume | | Pollen Viability | |
| | **Parent** | **$F_2$** | **Parent** | **$F_2$** | **Parent** | **$F_2$** |
| **Stripe** | 331 | 276 | 80 | 81 | 84 | 50 |
| **Non-stripe** | 183 | 125 | 1 | 27 | 1 | 18 |

| | *Fitness Traits* | |
|---|---|---|
| Foliar Phenotype | Seed Number & Average Seed Weight (per individual seed and per calyx) | |
| | **Parent** | **$F_2$** |
| **Stripe** | 54 | 56 |
| **Non-stripe** | 37 | 17 |

Greenhouse Experiment Results

**(Q1) Could foliar striping be part of a larger pollination syndrome (i.e., is it genetically correlated with traits known to affect pollinator attraction)?**

Corolla width did not differ significantly between foliar striping phenotypes or genotypes in the greenhouse experiments (Supplemental Table 2; Supplemental Figure 1A), even though parent plants produced slightly wider corollas than $F_2$ plants.

Corolla height was significantly affected by foliar striping phenotype, genotypes, and their interaction (Supplemental Table 2; Supplemental Figure 1B). Striped plants produced corollas 1.05x taller than non-striped plants. Parent plants produced corollas 1.03x taller than $F_2$ plants. The phenotype by genotype interaction, was driven by the striped parent plants that produced corollas ~1.08x taller than all other phenotype by genotype pairings.

Binary herkogamy, quantified as the presence of reverse herkogamy (anthers positioned above stigmas), showed a significant effect of phenotype (Supplemental Table 2; Supplemental Figure 2C). Non-stripe plants exhibited a higher probability of expressing reverse herkogamy than stripe plants, corresponding to approximately 2x higher odds of reverse herkogamy in non-stripe plants. Genotype did not significantly affect the probability of reverse herkogamy.

**(Q2) Is foliar striping associated with increased rates of pollination and/or pollination success, resulting in higher relative reproductive fitness in striped plants?**

In the greenhouse experiments, seed number was significantly affected by phenotype but not genotype or mating system treatment (Supplemental Table 2; Supplemental Figure 2A). Foliar striped plants produced 1.33x more seeds than non-striped plants.

Average individual seed weight was significantly affected by phenotype, genotype, mating system treatment, and the phenotype by genotype interaction (Supplemental Table 2; Supplemental Figure 2B). Non-stripe plants produced seeds that were 1.06x heavier than seed produced from striped plants. Parent plants produced seeds that were 1.14x heavier than $F_2$ family seeds. Outcrossed flowers produced seeds that were 1.26x heavier than seeds from selfed flowers. This phenotype by genotype interaction was largely driven by non-striped parent plants, as they produced seeds ~1.2x heavier than all other phenotype by genotype pairings.

Average seed weight per calyx (reproductive clutch) was significantly affected by phenotype and mating system treatment but not genotype (Supplemental Table 2; Supplemental Figure 2C). Striped plants produced reproductive clutches that were 1.25x heavier than the non-striped plants. Outcrossed flowers produced reproductive clutches that weighed 1.47x more than selfed flowers.

**Supplemental Table 2.** Results of analysis of variance for the greenhouse dataset demonstrating the effects of phenotype, genotype, and treatment (where appropriate) and their interaction on crimson monkeyflower traits. Any interactions not included here are due to their exclusion from the best fit models. Factors with significant effects are in bold type. Test-statistic is the F-statistic unless otherwise indicated.

| **Greenhouse Experiments** | | | | | |
|---|---|---|---|---|---|
| **Traits** | | **Fixed Effects** | **DF (numerator, denominator)** | **Test-statistic** | **p-value** |
| **Floral** | **Corolla Width (mm)** | Phenotype | 1, 184.31 | 0.097 | 0.756 |
| | | Genotype | 1, 228.14 | 3.034 | 0.083 |
| | **Corolla Height (mm)** | **Phenotype** | 1, 219.21 | 47.957 | **< 0.001** |

| | | | | | |
|---|---|---|---|---|---|
| | | **Genotype** | 1, 219.21 | 20.773 | **< 0.001** |
| | | **P*G Interaction** | 1, 219.21 | 22.147 | **< 0.001** |
| | **Herkogamy - Reverse vs. Approach** | **Phenotype** | 1 | $\chi^2$ = 11.574 | **< 0.001** |
| | | Genotype | 1 | $\chi^2$ = 0.821 | 0.365 |
| | **Nectar Volume (uL)** | Phenotype | 1, 186 | 0.0014 | 0.971 |
| | | **Genotype** | 1, 186 | 17.331 | **< 0.001** |
| | **Pollen Viability (%)** | **Phenotype** | 1 | $\chi^2$ = 34.169 | **< 0.001** |
| | | Genotype | 1 | $\chi^2$ = 2.498 | 0.114 |
| | | **P*G Interaction** | 1 | $\chi^2$ = 14.862 | **< 0.001** |
| **Fitness** | **Seed Number** | **Phenotype** | 1, 156 | 12.435 | **< 0.001** |
| | | Genotype | 1, 156 | 0.082 | 0.775 |
| | | Treatment | 1, 156 | 3.562 | 0.061 |
| | | P*G*T Interaction | 1, 156 | 0.572 | 0.451 |
| | **Average Seed Weight per Individual Seed** | **Phenotype** | 1, 159 | 3.901 | **< 0.05** |
| | | **Genotype** | 1, 159 | 23.328 | **< 0.001** |
| | | **Treatment** | 1, 159 | 90.426 | **< 0.001** |
| | | **P*G Interaction** | 1, 159 | 26.578 | **< 0.001** |
| | **Average Seed Weight per Calyx** | **Phenotype** | 1, 160 | 9.200 | **< 0.01** |
| | | Genotype | 1, 160 | 0.831 | 0.363 |
| | | **Treatment** | 1, 160 | 31.645 | **< 0.001** |
| **PCA - floral / nectar /** | **PC1** | Phenotype | 2, 45 | t = -0.21 | 0.83 |
| | | **Genotype** | 2, 45 | t = 2.75 | **< 0.01** |
| | **PC2** | Phenotype | 2, 45 | t = -0.176 | 0.86 |
| | | Genotype | 2, 45 | t = 0.59 | 0.56 |
| | **PC3** | Phenotype | 2, 45 | t = -0.12 | 0.90 |
| | | Genotype | 2, 45 | t = -0.23 | 0.82 |
| **PCA - floral / nectar** | **PC1** | Phenotype | 2, 186 | t = -1.28 | 0.20 |
| | | **Genotype** | 2, 186 | t = 4.18 | **< 0.001** |
| | **PC2** | Phenotype | 2, 186 | t = 1.43 | 0.15 |
| | | Genotype | 2, 186 | t = -1.24 | 0.22 |
| | **PC3** | Phenotype | 2, 186 | t = -0.77 | 0.44 |
| | | **Genotype** | 2, 186 | t = -4.41 | **< 0.001** |
| **P** | **PC1** | **Phenotype** | 2, 27 | t = -4.00 | **<0.001** |

| | | | | | |
|---|---|---|---|---|---|
| | | Genotype | 2, 27 | t = 1.02 | 0.32 |
| | **PC2** | Phenotype | 2, 27 | t = 0.74 | 0.47 |
| | | Genotype | 2, 27 | t = -1.52 | 0.14 |
| | **PC3** | Phenotype | 2, 27 | t = -0.44 | 0.67 |
| | | Genotype | 2, 27 | t = -0.92 | 0.37 |

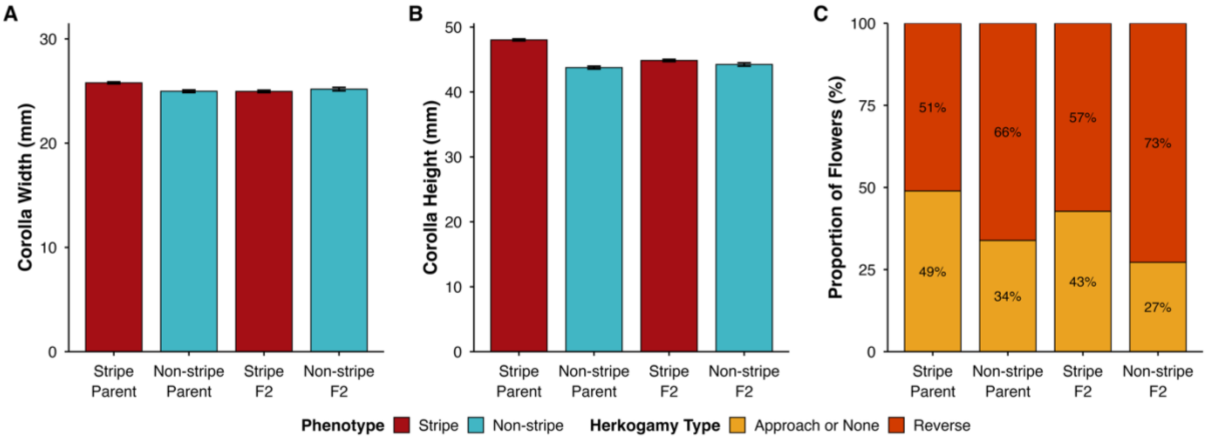

**Supplemental Figure 1.** Floral traits of Parent and $F_2$ plants with and without foliar striping; error bars in panels A and B represent +/- 1 SE from the mean. (**A**) Corolla width, (**B**) Corolla height, and (**C**) Binary herkogamy are from the greenhouse datasets.

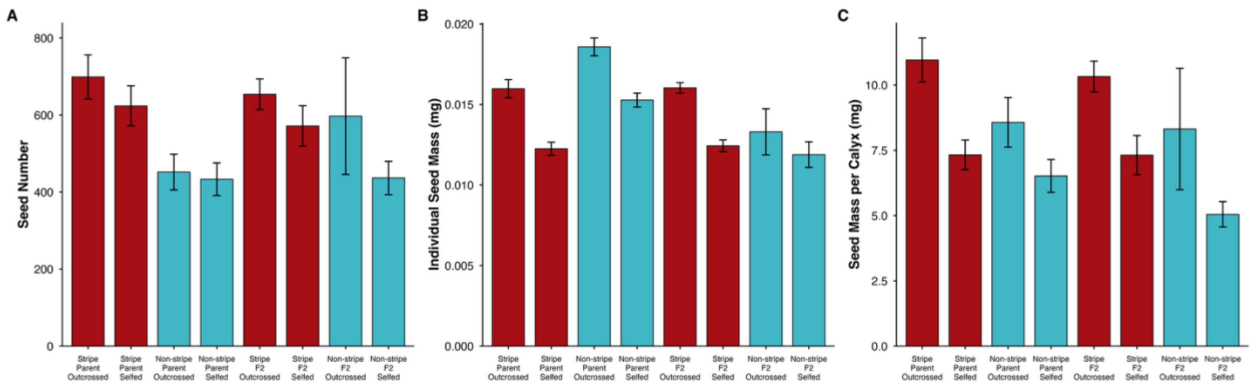

**Supplemental Figure 2.** Seed set traits of plants with and without foliar striping, considering genotype and mating system treatment. All data is from the greenhouse experiment plants. Error bars represent +/- 1 SE from the mean. (**A**) Seed Number for all plants that produced at least one seed, (**B**) Average seed weight (mg), (**C**) Average seed weight for all seeds in a calyx (mg).

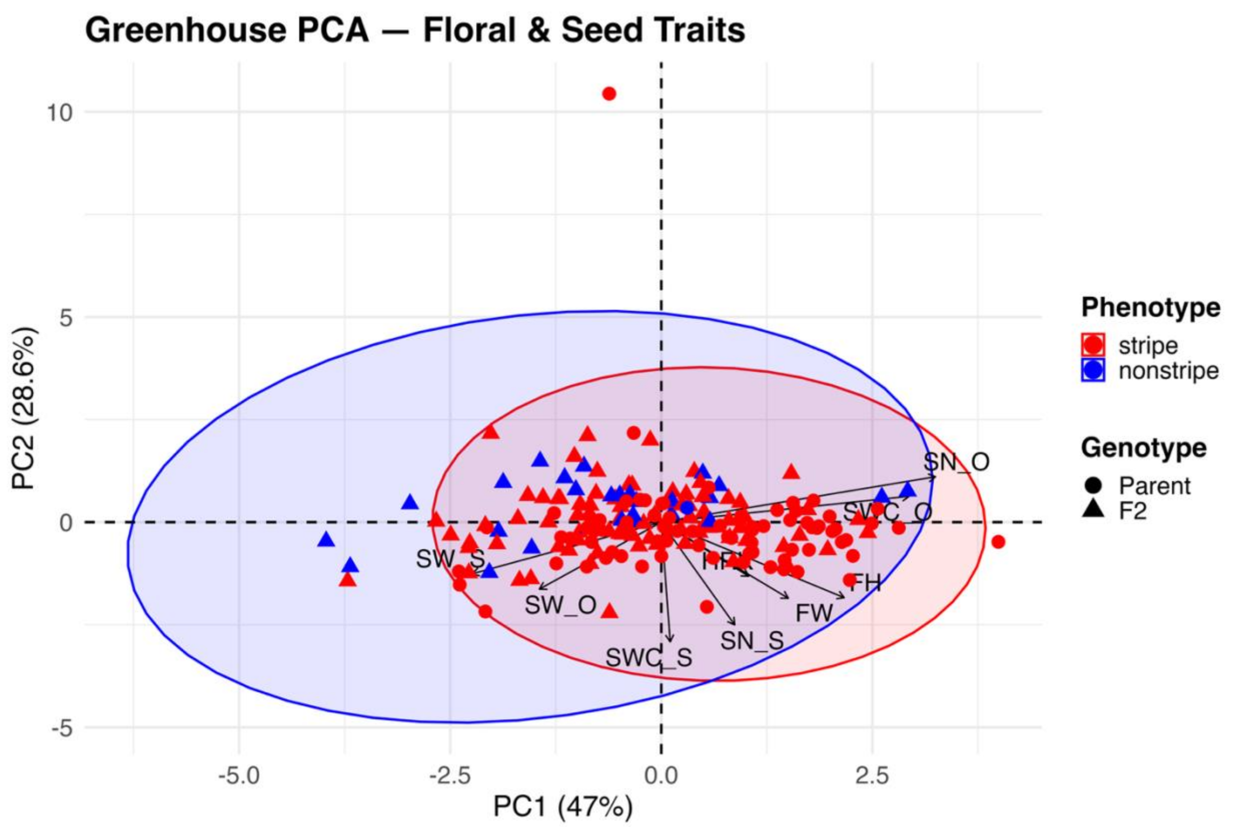

**Supplemental Figure 3.** Combined floral and seed trait PCA from greenhouse data. Phenotype and genotype are visually represented, though not included as part of the PC axes. Vector abbreviations are: (FW) flower width, (FH) flower height, (HRK) herkogamy, (SN_O) seed number-outcrossed, (SN_S) seed number-selfed, (SW_O) seed weight-outcrossed, (SW_S) seed weight-selfed, (SWC_O) seed weight calyx-outcrossed, (SWC_S) seed weight calyx-selfed.

<u>Additional lines of thoughts on foliar striping that were not pursued beyond pilot studies:</u>

Prior to hypotheses proposed in this publication, we considered additional hypotheses about the importance of the anthocyanin stripe. Preliminary information on chemical defenses and the UV-spectrum has been included below.

To determine if the anthocyanins are additionally visible on the UV spectrum available to hummingbirds, we collaborated with Dr. Kelsey Byers (The John Innes Centre, Norwich Research Park). Results from UV photography and reflectance spectroscopy of the crimson monkeyflower leaves from the striped (MvBL) line, show no UV reflectance in either the green leaves or the stripe, and no UV patterning. The data are from three separate leaves (same leaves for both UV photography and reflectance spec).

(A)

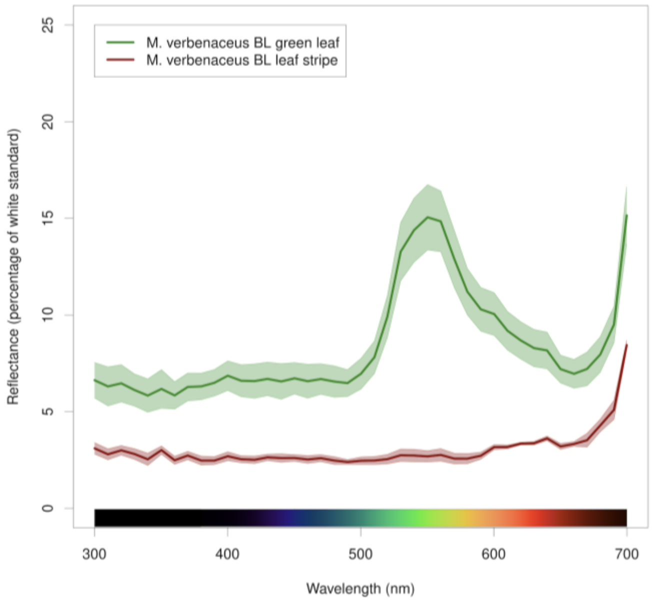

(B)

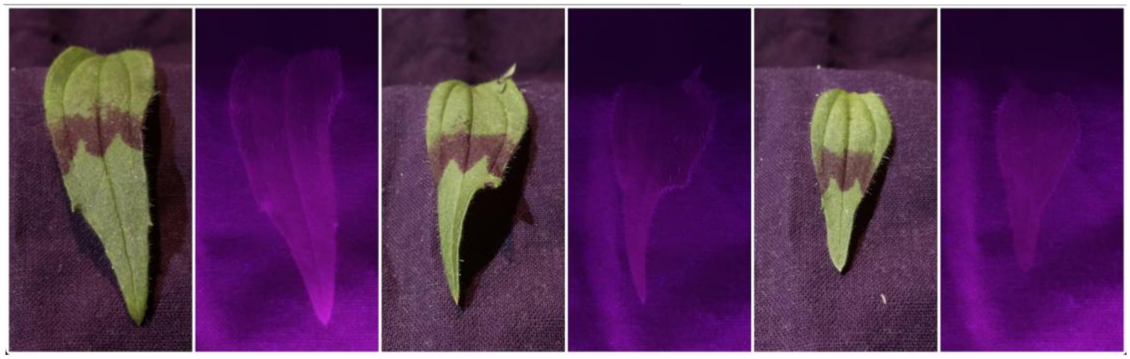

**Supplemental Figure 4.** (**A**) Reflectance spectroscopy, and (**B**) UV photography of foliar striping on crimson monkeyflowers.
